# Identification of an inhibitory pocket in falcilysin provides a new avenue for malaria drug development

**DOI:** 10.1101/2021.04.08.438947

**Authors:** Grennady Wirjanata, Jianqing Lin, Jerzy Michal Dziekan, Abbas El Sahili, Zara Chung, Seth Tjia, Nur Elyza Binte Zulkifli, Josephine Boentoro, Roy Tham, Lai Si Jia, Ka Diam Go, Han Yu, Anthony Partridge, David Olsen, Nayana Prabhu, Radoslaw M Sobota, Pär Nordlund, Julien Lescar, Zbynek Bozdech

**Affiliations:** School of Biological Sciences, Nanyang Technology University, Singapore 637551; NTU Institute of Structural Biology, Nanyang Technology University, Singapore 637551; Infectious Diseases Labs & Singapore Immunology Network, Agency for Science, Technology and Research, 138648, Singapore; MSD, Singapore; Merck & Co., Inc., West Point, PA 19486 USA; Institute of Molecular and Cell Biology, Agency for Science, Technology, and Research (A*STAR), Singapore 138673; Functional Proteomics Laboratory, Institute of Molecular and Cell Biology, Agency for Science, Technology and Research (A*STAR), Singapore, Singapore; Department of Oncology and Pathology, Karolinska Institutet, Stockholm, 17177 Sweden; Antimicrobial Resistance Interdisciplinary Research Group, Singapore-MIT Alliance for Research and Technology, Singapore 637551

**Author notes:** Equally contributing authors (Grennady Wirjanata, Jianqing Lin).

## Abstract

Despite their widespread use, our understanding of how many antiparasitic drugs work remains limited. We used mass-spectrometry based cellular thermal shift assay (MS-CETSA) to identify possible protein targets of several malaria drugs and drug candidates. We found that falcilysin (FLN) is a common target for several quinoline drugs including chloroquine and mefloquine, as well as drug candidates MK-4815, MMV000848 and MMV665806. At pH 7.5, these compounds all inhibit FLN proteolytic activity with IC_50_ values ranging from 1.6 to 67.9 µM. Their interaction with FLN was systematically probed by isothermal titration calorimetry and X-ray crystallography, revealing a shared hydrophobic pocket in the catalytic chamber of the enzyme. Characterization of transgenic cell lines with depleted FLN expression demonstrated statistically significant increases in susceptibility towards chloroquine, mefloquine, MK-4815 and MMV000848. Taken together, our findings point to a multimodal mechanism of action for several commonly used anti-malaria drugs. Importantly, a common allosteric pocket of FLN appears amenable to inhibition, providing a structural basis to guide the development of novel drugs against malaria.

## Introduction

With more than 240 million cases and 600,000 deaths in 2020, malaria remains a major public health problem (World Health Organization, 2021). Declining efficacy for artemisinin-based combination therapy (ACT) by emerging resistance to ACT components in Southeast Asia and Africa is particularly worrying, given the limited number of clinically approved treatments (Ashley et al., 2014; Balikagala et al., 2021; Dondorp et al., 2009; Phyo et al., 2016; van der Pluijm et al., 2019). Novel drugs are urgently needed not only to counter the spread of drug resistance but also to combat malaria on the world-wide scale in the upcoming decades. Antimalarial drugs, including those in current clinical use, were discovered largely through phenotypic screening using cell-based assays to determine their efficacy for inhibiting parasite growth (Hovlid and Winzeler, 2016). These include artemisinin-as well as quinoline-based compounds that currently constitute the backbone of the ACTs, the frontline treatment of malaria worldwide (Su and Miller, 2015; van Schalkwyk). Given their historical success, cell-based phenotypic screens were also employed to identify novel lead candidates derived from chemical libraries including the GlaxoSmithKline Tres Cantos Antimalarial Set (TCAMS) and Medicines for Malaria Venture (MMV) Malaria Box (Gamo et al., 2010; Spangenberg et al., 2013). The main drawback of the phenotypic screens is their agnostic approaches in which, initially, nothing is known about the biological activity of the selected compounds. As such there is an immediate need to study the mechanism of action (MOA) of the compounds to develop these chemical scaffolds into clinically relevant drugs (Challis et al., 2022).

To address this challenge, several anti-malaria drug target identification methods have been developed in the last two decades. One of the main methods involves *in vitro* selection of resistance to a compound coupled with whole genome sequencing (IVIEWGA) (Baragana et al., 2015; Cowell et al., 2018; Favuzza et al., 2020; Kuhen et al., 2014; Luth et al., 2018; Rottmann et al., 2010; Yang et al., 2021). The mutations resulting from these selections are good indicators of proteins and/or pathways affected by the drugs and as such, provide important additional information regarding their MOA. This method has been successful in identifying or reconfirming antimalarial drug targets, such as the proteasome (Xie et al., 2018), thymidylate synthase, farnesyltransferase, dipeptidyl aminopeptidase 1, aminophospholipid-transporting P-type ATPase (Cowell et al., 2018), protein kinase CLK3 (Alam et al., 2019), acetyl-coenzyme A synthetase (Summers et al., 2022) and isoleucil tRNA synthetase (Istvan et al., 2023). Upon identification of such genes, complementary studies are typically conducted to discern the actual drug/compound targets from factors of resistance (Luth et al., 2018). Such studies include metabolic profiling of drug-treated parasites (Alam et al., 2019; Allman et al., 2016; Cowell et al., 2018; de Vries et al., 2022; Gisselberg et al., 2018; Istvan et al., 2023; Murithi et al., 2020; Summers et al., 2022), modelling and crystallography studies (Istvan et al., 2023; Summers et al., 2022; Xie et al., 2022), and CRISPR-based allelic replacement studies (Istvan et al., 2023; Summers et al., 2022; Xie et al., 2022). In the case where resistant parasites cannot be selected using IVIEWGA, chemoproteomics approaches are typically considered. This includes approaches such as affinity-(Paquet et al., 2017) and clickable-based probes (Ismail et al., 2016; Wang et al., 2015) and protein stability-based target identification approaches such as Stability of Proteins from Rates of Oxidation (SPROX) (Lu et al., 2020) or Mass Spectroscopy-linked Cellular Thermal Shift Assay (MS-CETSA) (Dziekan et al., 2020b; Dziekan et al., 2019). The latter has emerged recently as a powerful tool to study drug target engagement for a variety of drug target discoveries ranging from anti-cancer (Jafari et al., 2014; Martinez Molina et al., 2013; Martinez Molina and Nordlund, 2016), to antimicrobials of human pathogens, such as *E. coli* (Hart et al., 2019), *Toxoplasma gondii* (Herneisen and Lourido, 2021), *Leishmania donovani* (Chakrabarti et al., 2022), and *P. falciparum* (Dziekan et al., 2019; Lu et al., 2020; Milne et al., 2022).

The initial rationale of the present study was to utilize MS-CETSA to identify protein interacting partners of quinoline-based antimalarial drugs that represent one of the most important classes of antimalarials. Here we hypothesized that these proteins may represent new chemotherapeutic targets for future antimalaria drug discovery and development. Here we uncovered distinct sets of proteins that may constitute or at least contribute to quinoline MOA, from which falcilysin (FLN) was widely shared across the drugs. Coincidently, MS-CETSA revealed interaction between FLN and two additional experimental compounds from the MMV box: MMV665806 and MMV000848, and one pre-clinical lead compound from Merck: MK-4815 (Powles et al., 2012). Using X-ray crystallography, we identified a hydrophobic pocket in the catalytic chamber of FLN as a common binding site for all these compounds. Overall, our work introduces a “druggable” pocket in FLN that deserves further studies for the development of antimalaria drugs.

## Results

### MS-CETSA of quinoline antimalarials identifies FLN as a key protein target

Here we used MS-CETSA to interrogate the MOA of several quinoline-based ACT partner drugs including amodiaquine (AQ), pyronaridine (PYD), piperaquine (PIP) and lumefantrine (LUM). In addition, we included the chemically related chloroquine (CQ), historically the most successful quinoline-based antimalaria chemotherapeutic. An identical strategy previously led to the discovery of PfPNP as a direct interacting partner of quinine (Q) and mefloquine (MFQ) (Dziekan et al., 2020b; Dziekan et al., 2019). Specifically, we carried out the Iso-Thermal Dose Response (ITDR) implementation of MS-CETSA within a range of drug concentrations covering both their *in vitro* IC_50_ values and their physiological therapeutic concentrations (∼1 nM to 10 µM). The initial set of ITDR measurements was carried out with *P. falciparum* live trophozoites (*in-cell* settings). In these settings, the thermal shifts can represent direct protein-drug binding but also other drug-induced effects, such as disruptions of protein-protein interactions, metabolite binding or posttranslational modifications; all related to the drug’s MOA (Dziekan et al., 2020b; Dziekan et al., 2019). Indeed, the *in-cell* ITDR uncovered distinct sets of proteins for each tested drug **(Fig. 1A)**. While LUM and PIP each only caused thermal shift of a single protein, AQ, CQ, and PYR thermostabilized four, nine and sixty-six proteins, respectively (**Table S1**). Even though each drug appeared to have largely unique effects, there were several proteins shared across at least two drugs including *cysteine--tRNA ligase* (PF3D7_1015200), *asparagine and aspartate rich protein 1* (PF3D7_1233600), *CUGBP Elav-like family member 1* (PF3D7_1359400) and *Hsp70/Hsp90 organizing protein* (PF3D7_1434300). Overall, our result is compatible with the current view of the MOAs of quinoline drugs that are expected to be pleiotropic, in which each drug acts through a distinct set of molecular targets in the *Plasmodium* cell. Nevertheless, we also identified the metalloprotease *falcilysin* (FLN, PF3D7_1360800) that was thermostabilized by four out of the five quinolines possibly representing a common quinoline target (**Fig. 1A**).

**Figure 1.**
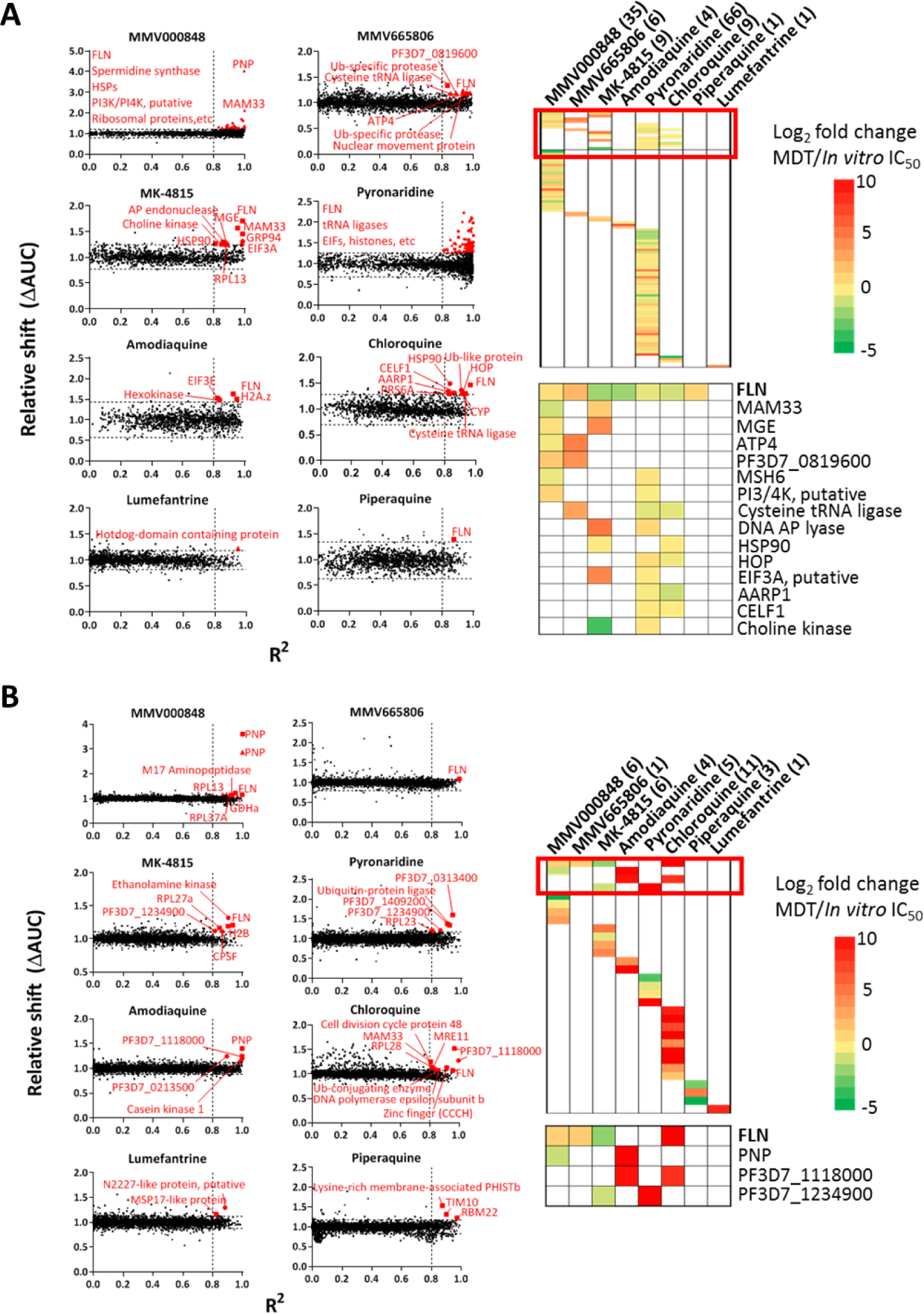
Protein Hits identified by MS-CETSA. Scatter plots on the left side of the figures represent distribution of protein stabilization obtained from in-cell (A) and lysate (B) MS-CETSA, plotted as a function of R^2^ value (goodness of curve fit) against ΔAUC (area under the curve of heat-challenged sample normalized against non-denaturing 37°C control) for all proteins detected in the assay. Two and a half of median absolute deviation (MAD) of ΔAUC in each dataset (MAD × 2.5) and R^2^ = 0.8 cutoffs are indicated on the graph. Proteins with significant thermal stabilization are labeled. Comprehensive list of protein hits with their respective information is available in Table S1. Heatmaps on the right side of the figures represent the overview of protein hits by all compounds (top) and zoomed in list of common protein targets identified in multiple compounds (bottom). Bracketed numbers after compound name represent the number of stabilized proteins by each compound. Color gradient represents log2 fold change ratio of minimal dose threshold (MDT) against *in vitro* growth IC_50_ of each compound. For comparison purpose, the following *in vitro* growth IC_50_ values are used for the calculation: MMV000848=500 nM, MMV665806=400 nM, MK-4815=400 nM, amodiaquine=35 nM, pyronaridine=30 nM, chloroquine=25 nM, piperaquine=40 nM, and lumefantrine=20 nM. All hits were identified from a total of two independent experiments for chloroquine, MK-4815, amodiaquine, piperaquine, and lumefantrine, whereas this from MMV665806, MMV000848, and pyronaridine were identified from a single experiment.

To explore this further, we identified three additional compounds that exhibit strong antimalarial activities and also interact with FLN. These include two experimental compounds from the MMV malaria box (MMV000848 and MMV665806) and the pre-clinical antimalaria compound MK-4815 (Powles et al., 2012) (**Fig. 1A and Table S1**). In an identical *in-cell* ITDR assay, the three compounds also affected the thermostability of FLN and other largely distinct sets of proteins: six (6) for MMV665806, nine (9) for MK-4815 and 33 for MMV000848 (**Fig. S2**). Likewise, there were some overlaps between the proteins stabilized by the three compounds and quinolines. These included *DNA-apurinic lyase* (PF3D7_0305600), *heat shock protein 90* (PF3D7_0708400), *eukaryotic translation initiation factor 3 subunit A* (PF3D7_1212700) and *choline kinase* (PF3D7_1401800) thermostabilized by MK-4815 and *DNA mismatch repair protein MSH6* (PF3D7_0505500) and *phosphatidylinositol 3- and 4-kinase* (PF3D7_0311300) thermostabilized by MMV00084, respectively, and at least one quinoline drug (**Fig. 1A**). This suggests some overlap between the MOA quinolines and the experimental compounds that reaches beyond FLN binding. The relevance of the measured thermal shifts to the drug/compound MAOs is also demonstrated by good correlations between the ITDR minimal dose threshold (MDT, minimum dose needed to induce thermal stabilization) and their *in vitro* parasiticidal activity (IC_50;_ 50% inhibitory concentration) (**Table S1, Fig. 1A**, *right panel*s). Indeed, most identified thermal shifts correlated well with the growth inhibitory activity, with median MDT/IC_50_ ratio ranging from 0.82 to 3 for MMV000848, MK-4815, CQ, PYD, AQ, and PIP. In particular, we observed low MDT/IC_50_ ratios (0.11 to 2) for FLN engagement with MK4815 and AQ further suggesting a high affinity of this interactions with high probability of taking part in the drug/compound MOA. It is noteworthy that both MMV compounds thermostabilized *non-SERCA-type Ca2+-transporting P-ATPase* (PF3D7_1211900, PfATP4) that is currently being pursued as a potential drug target (Flannery et al., 2015; Rottmann et al., 2010; Spillman et al., 2013).

To complement the *in-cell* MS-CETSA, we carried out lysate-based ITDR measurements that are more reflective of drug-target interaction (Dziekan et al., 2020b; Dziekan et al., 2019). Indeed, the lysate-based ITDR identified between one to eleven proteins thermostabilized by the eight compounds (**Fig. 1B**), with only limited overlap with the *in-cell* results (**Fig. 1A and Table S2**). Crucially, like in the *in-cell* conditions, the lysate MS-CETSA identified FLN as the most frequent hit, being stabilized by CQ, MK-4815, MMV000848 and MMV665806. Further inspection of the FLN ITDR stabilization profile indicates good agreement between the MDT values generated by lysate and *in-cell* ITDR for the three experimental drugs, ranging from ∼100 nM in MK-4815 through ∼1 µM in MMV000848 and MMV665806. This contrasts CQ, with MDT values of 14 nM and 56.1 µM in *in-cell* and lysate experiments, respectively (**Fig S1**). We observed no thermostabilization of FLN in the lysate ITDR with AQ, PIP, PYR, which suggests that these drugs require a specific cellular environment to interact with FLN. Curiously, we observed AQ and MMV000848-induced stabilization of PfPNP that was previously shown to interact with two other quinoline drugs such as quinine and mefloquine (MFQ) (Dziekan et al., 2020b; Dziekan et al., 2019). Taken together, the lysate ITDR results support our main *in-cell* observation of FLN being an important target in quinoline or quinoline-like antimalaria drugs and as such could serve as a potential target. In the following parts of this study, we carried biochemical, structural, and transgenic studies to evaluate this hypothesis.

### Quinolines, MK-4815, MMV000848 and MMV665806 bind to FLN and inhibit its proteolytic activity

First, we conducted isothermal titration calorimetry (ITC) measurements with a recombinant FLN enzyme expressed heterologously in *E. coli* (**Fig. 2**). The ITC results showed a 1:1 stoichiometry in FLN binding for the four compounds that caused a thermal shift in the lysate ITDR assay (**Fig. 1B)**. Out of these, MK-4815 and MMV000848 bind with the highest affinities with dissociation constants K_d_ of 1.79 ± 0.306 µM and 1.98 ± 0.183 µM, respectively, followed by MMV665806 (K_d_ = 7.33 ± 2.79 µM) (**Fig. 2A**). In addition, the ITC results confirmed that FLN interacts with CQ (K_d_ =116 ± 18.8 µM) and MFQ (K_d_ =26.7 ± 8.6 µM) which was previously shown to induce a thermal shift of FLN (Dziekan et al., 2019). Due to the low affinity, the CQ-FLN interaction was determined by a displacement assay rather than a direct ITC assay (see methods) (**Fig. 2A)**. Furthermore, to confirm the binding specificity of FLN, we performed a negative control whereby FLN was titrated with artemether, a chemically unrelated drug, and no binding was observed (**Fig. S2B)**.

**Figure 2.**
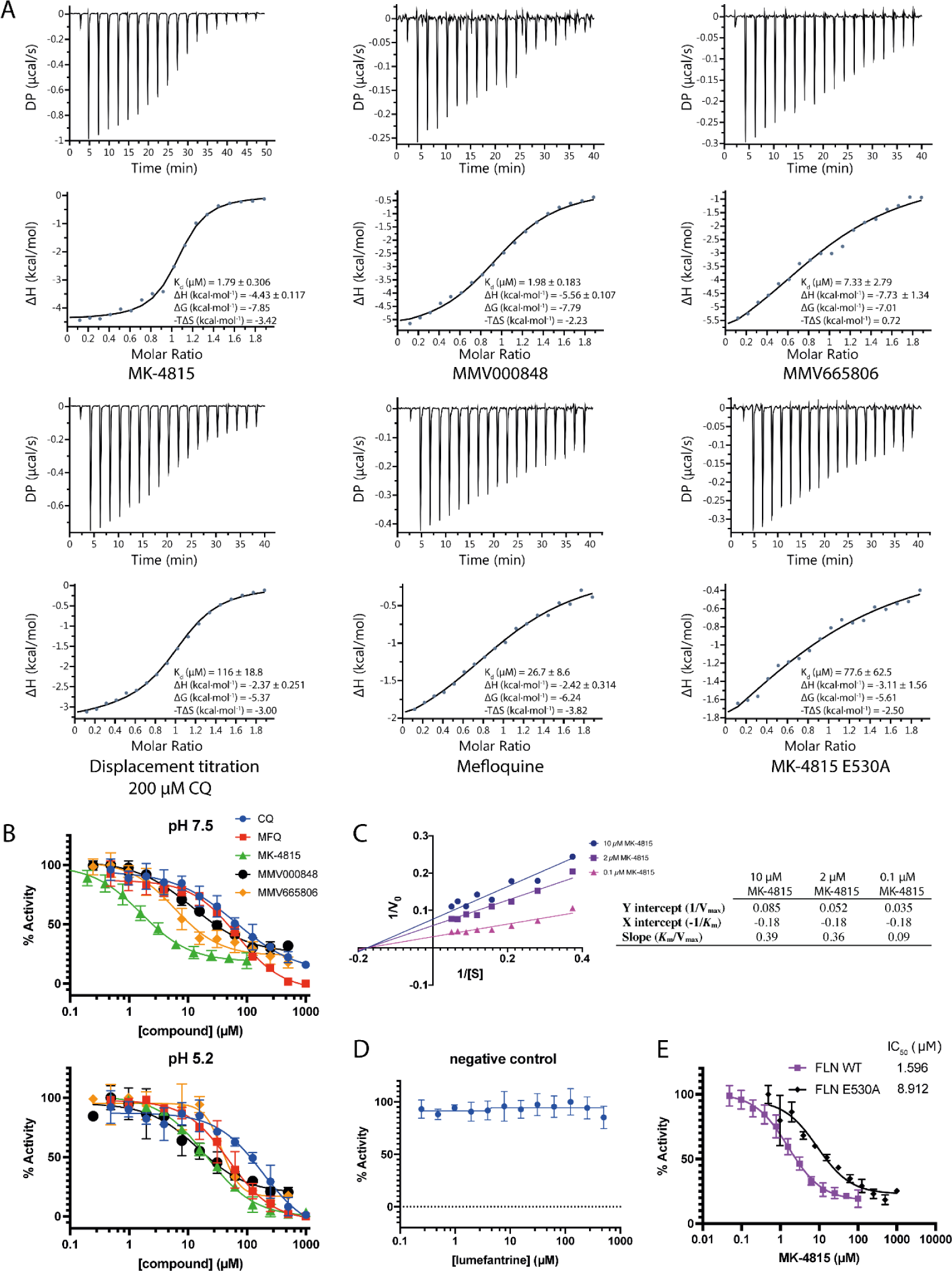
FLN binding and inhibition by CQ, MFQ, MK-4815, MMV000848 and MMV665806. **(A)** Isothermal titration calorimetry measurement between FLN and compounds. **(B-E)** FLN enzymatic inhibition assay. Protease activities were measured in triplicates and normalized for each compound. Error bars denote standard deviations. **(B)** FLN is inhibited by all five compounds. IC_50_ values are summarized in Table 1. **(C)** Double-reciprocal (Lineweaver-Burk) plot of FLN enzymatic activity in the presence of MK-4815. K_m_ is unchanged while V_max_ decreases as MK-4815 concentration increases, suggesting non-competitive inhibition. **(D)** Lumefantrine was used as a negative control for FLN enzymatic inhibition assay. No inhibition was observed. **(E)** Effect of E530A mutation on FLN enzymatic inhibition by MK-4815 at pH 7.5. IC_50_ increases from 1.6 μM to 8.9 μM because E530A mutation weakens MK-4815 binding (see Fig. 3E).

Next, we carried out enzymatic inhibition activity assays using a ten-amino-acid peptide substrate at pH 7.5 and 5.2, reflecting putative conditions in plastid/mitochondria and DV, respectively (see below) (**Fig. 2B**). Measurements listed in **Table 1** reveal a range of IC_50_ values with the strongest inhibition found at pH 7.5 for MK-4815 (IC_50_=1.611 µM), MMV665806 (IC_50_ = 5.943 µM) and MMV000848 (IC_50_ = 11.86 µM). Lumefantrine that did not induce any thermal shift (**Fig. 1**), presumably not interacting with FLN, failed to inhibit its proteolytic activity (**Fig. 2D**) and as such was used as a negative control for the FLN inhibition assay. The two quinolines, CQ and MFQ exhibited somewhat weaker inhibition with respective IC_50_ values of 63.81 µM and 67.94 µM. At pH 5.2, the inhibition potential of most studied compounds dropped to some degree, ranging from MMV000848 (IC_50_ of 14.74 µM ∼ 1.2-fold decrease) to CQ (IC_50_ of 186.2 µM ∼ 3-fold decrease) to MK-4815 (IC_50_ of 20.58 µM ∼50-fold decrease). A lower pH appears to only enhance the inhibition by MFQ, with IC_50_ of 49.26 µM which represents a 1.4-fold improvement compared to the inhibition at neutral pH. Finally, to decipher the mode of enzymatic inhibition, we utilized the most potent FLN inhibitor MK-4815 at pH 7.5 and carried out a Lineweaver-Burke analysis (**Fig. 2C**). With increasing concentrations of MK-4815, the value of V_max_ decreases whilst the K_m_ remains unchanged, suggesting noncompetitive inhibition of FLN protease activity. These kinetics measurements, where inhibitor binds both free enzyme and enzyme-substrate complex, point to the presence of an allosteric pocket in FLN accounting for enzymatic inhibition. Crucially, for each of the five compounds, the K_d_ values are comparable to the IC_50_ values derived from FLN enzymatic inhibition (**Table 1**). For MK-4815, MMV000848 and MMV665806, this correlation extends to antiparasitic IC_50_ values measured in cell-based assays which exhibit by comparable values in the low µM range that are consistent with the biochemical data. CQ and mefloquine appear as notable exceptions with *in vitro* anti-parasitic IC_50_ values of 0.025 µM and 0.030 µM, three orders of magnitude lower than their FLN enzymatic inhibition, suggesting a complex MOA.

**Table 1.** Comparison of values for MDT, dissociation constants K_d_ (Fig. 2A), FLN proteolytic inhibition IC_50_ (Fig. 2B) and in vitro parasite growth inhibition IC_50_.

|  | Chloroquine | Mefloquine | MK-4815 | MMV000848 | MMV665806 |
| --- | --- | --- | --- | --- | --- |
| MDT (μM) | 56.10 | 1 μM<sup>#</sup> | 0.09 | 1.10 | 1.03 |
| K<sub>d</sub> (μM) | 116 ± 18.8 | 26.7 ± 8.6 | 1.79 ± 0.30 | 1.98 ± 0.18 | 7.33 ± 2.79 |
| FLN inhibition |  |  |  |  |  |
| IC<sub>50</sub> pH 7.5 (μM) | 63.81 | 67.94 | 1.61 | 11.86 | 5.94 |
| 95% CI | 36.84-338.20 | 53.26-92.99 | 1.20-2.11 | 8.82-16.35 | 3.97-8.51 |
| IC<sub>50</sub> pH 5.2 (μM) | 186.2 | 49.26 | 20.58 | 14.74 | 37.62 |
| 95% CI | 113.3-1344.0 | 37.98-66.45 | 17.81-23.74 | 8.10-35.14 | 30.17-48.14 |
| Anti-parasitic |  |  |  |  |  |
| *In vitro* IC<sub>50</sub> (μM) | 0.025 | 0.030 | 0.400 | 0.500 | 0.400 |

<sup>#</sup> Intact cell assay

### Crystal structures of FLN bound to anti-malaria compounds

To better understand the structural basis for FLN inhibition, we obtained high resolution co-crystal structures of FLN bound to CQ, MFQ, MK-4815, MMV000848 and MMV665806 at 1.82, 1.55, 1.9, 1.96 and 2.0 Å resolution respectively (**Fig. 3 and Table S3**). In the closed conformation of FLN that was captured, the large reaction chamber of the active site occupies a volume of 10,886 Å^3^ which is fully enclosed by the N and C terminal halves of the protein. A general view of FLN is given in **Fig. 3A** highlighting the topography of its peptide substrate binding cavity and the spatial distance between the shared inhibitor binding site and the M16C protease active site. The protease active site of FLN contains a catalytic zinc ion tetrahedrally coordinated by His129 and His133, which belong to the inverted 129-HXXEH-133 motif, Glu243 and also by a water molecule (**Fig. 3B)**. For all five compounds (listed above), clear residual electron density at a distance of ∼24 Å away from the metalloprotease active site was seen in a hydrophobic pocket housed between helices α13 and α14 of the enzyme. This pocket is lined by residues Phe82, Phe527, Ile 544 and Phe545 of FLN that form a hydrophobic patch largely exposed at the surface of the catalytic chamber (**Fig. 3**).

**Figure 3.**
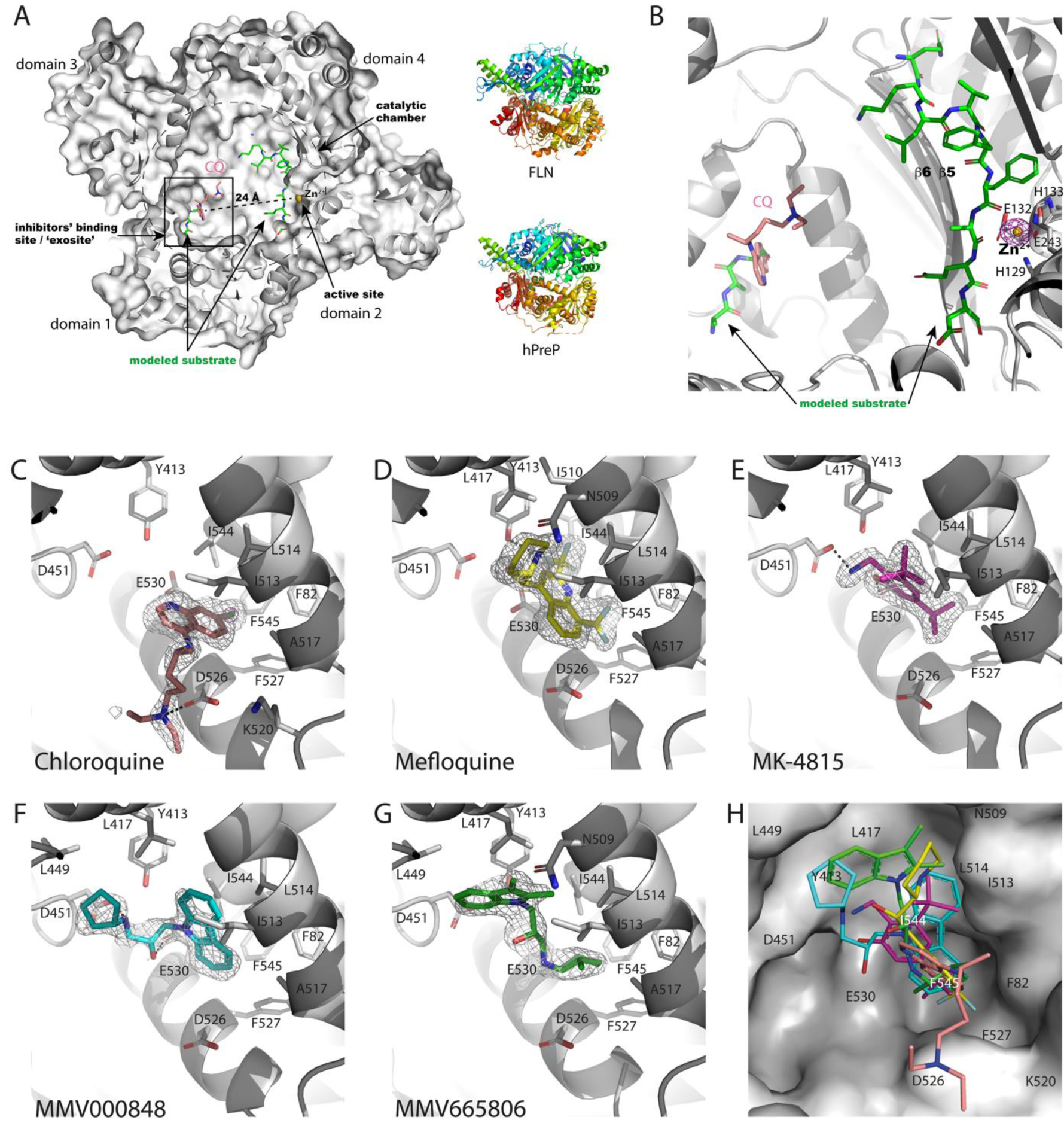
Crystal structures of FLN bound to inhibitors. **(A)** Sliced view of the FLN catalytic chamber (delineated by a dashed circle) in complex with CQ (pink sticks). The catalytic zinc ion is shown as a gold sphere. The inhibitors’ binding pocket (boxed) is approximately 24 Å from the catalytic zinc. To generate this figure, another M16C metalloprotease hPreP in complex with its peptide substrate (PDB access code 4NGE) was superimposed to FLN to generate a FLN-substrate model. hPreP has 23% amino-acid sequence identity and a r.m.s.d. of 1.9 Å over 742 Cα superimposed atoms with FLN. The hPreP peptide substrate is displayed as green sticks while hPreP is omitted for clarity (left panel). Note that the inhibitors’ binding site overlaps with part of the hPreP peptide substrate binding site (previously reported as ‘exosite’). On the right panel, a rainbow representation of FLN and hPreP (PDB: 4NGE) shows high structural similarity. **(B)** Close-up view of the FLN catalytic chamber. FLN is shown as cartoon. Zinc-coordinating residues H129, H133 and E243 at the active site are displayed as sticks. An anomalous Fourier map collected at the Zinc absorption edge is shown as a purple mesh an contoured at a level of 8 σ. The modeled substrate forms an antiparallel β-sheet with strand β5 of FLN. Co-localization of inhibitors’ binding site and exosite indicates a possible steric hindrance between the inhibitors and long peptide substrates. **(C-G)** The inhibitors’ binding pocket of FLN where each inhibitor is shown as sticks. FLN is shown in grey. Simulated annealing F_0_-F_c_ omit Fourier maps are contoured at 2.5 σ and displayed as mesh. Salt bridges and hydrogen bonds are depicted as dotted lines. The omit map of CQ at 9 σ (purple mesh) confirms the position of Chloride atom which forms halogen-pi electron interaction with F545. (H) Superposition of inhibitors with a surface view of the binding pocket.

How do the various compounds interact with FLN using the same pocket? The chloride moiety of CQ points inwards establishing a halogen bond with the phenyl ring of Phe545, a residue strictly conserved in other *Plasmodium* FLN proteins such as *P. vivax* and *P. malariae* (**Fig. S3**). Parameters defining this interaction fall within the range of tabulated halogen-pi electron (C-Cl ---π) interactions (Matter et al., 2009): (i) the distance between the Cl atom and the centroid of the Phe545 phenyl ring is 4.1 Å, (ii) the angle between the CQ and Phe545 aromatic rings is 69.9° and (iii) the distance between the two rings centroids is 7.1 Å. Thus, this halogen-pi electron interaction plays a key role to orient and stabilize CQ inside the FLN catalytic chamber (**Fig. 3C**). Several aliphatic and aromatic residues Ile544, Leu514, Ile513, Phe82, Ala517 and Phe527 further contribute to CQ-FLN binding via van der Waals and hydrophobic interactions. In addition, the tertiary amine of CQ which is protonated at both acidic and neutral pH forms a salt bridge with Asp526 (**Fig. 3C**). However, despite its favorable enthalpic contribution, this interaction could be entropically unfavorable because it rigidifies the flexible tail of CQ. (**Fig. 3C**).

Likewise, MFQ is also mainly stabilized via hydrophobic interactions with Ile544, Leu514, Ile513, Phe82, Ala517, Phe527, Phe545, Leu417 and Ile510. Its six fluorine atoms are buried in the hydrophobic binding site except one fluorine that might form a weak hydrogen bond with Tyr413 (**Fig. 3D**). MK-4815 has one of its trimethyl group oriented towards Phe82, Phe527 and Phe545 while its other trimethyl group is surrounded by Leu417, Ile544, Leu514 and Ile513. In addition, its hydroxyl group hydrogen-bonds with Glu530 and its protonated amine group forms a salt bridge with Asp451. In an attempt to correlate the structural and biochemical observations, we mutated Glu530 to alanine (FLN-E530A mutant). The FLN-E530A single mutant showed much weaker binding to MK-4815 with a K_d_ of 77.6 μM (instead of K_d_ of 1.79 ± 0.306 µM for the WT enzyme) as measured by ITC (**Fig. 2A**). Consistently, the E530A mutation also decreases the potency of MK-4815 on FLN: enzymatic IC_50_ of MK-4815 increases from 1.6 µM on FLN WT to 8.9 µM on E530A mutant (**Fig. 2E**).

Both MMV000848 and MMV665806 mostly bind via hydrophobic interactions with Leu449, Leu417, Ile544, Leu514, Ile513, Phe545, Phe82 and Phe527. In addition, like in the case of MK-4815, the hydroxyl group of MMV000848 makes an hydrogen-bond with Glu530 and its protonated secondary amine group forms a salt bridge with Asp451 (**Fig. 3F,G**), highlighting a structural mimicry between the two Malaria box compounds and MK-4815. In summary, we identified an exposed hydrophobic pocket in the catalytic chamber of FLN, which constitutes a promiscuous binding site for quinolines, MMV000848, MMV665806 and MK-4815 with a clear impact of compound binding on enzyme activity (**Fig. 3H**).

### A possible mechanism of inhibition of FLN

What is the mechanism underlying the inhibition of FLN protease activity by the five compounds binding at a hydrophobic pocket ∼24 Å away from the metalloprotease active site? The Lineweaver-Burke kinetics analysis suggests an allosteric mode of inhibition in which inhibitor binding at a site distant from the active site induces a strain in the enzyme dynamic. FLN is a globular protein with a large reaction chamber centrally located between the four protein domains (**Fig. 3A**). In the present co-crystal structures with inhibitors, only the closed conformations of FLN was captured, with an overall conformation similar to other M16C metalloproteases such as the human presequence protease (hPreP) (King et al., 2014). However, in solution, a transition between closed and open conformation must occur to allow substrate entry. While the closed conformation is conducive to peptide cleavage, at the end of the cycle, FLN must reopen to release cleaved peptides. Structural evidence for such a cycle between open and closed forms was first shown experimentally for the human presequence protease, using small angle X-ray scattering (King et al., 2014). More recently, Liang and colleagues used CryoEM to trap the human presequence protease in an open form, providing high resolution information about the elusive open form and giving strong support for a switch between the closed and open forms during the proteolytic cycle of M16C metalloprotease in general (Liang et al., 2022). We also determined via cryo-EM a high resolution structure of FLN in an open conformation giving further support to this hypothesis of a dynamic transition (Lin J., Bozdech Z & Lescar, J manuscript in preparation). Taken together, an attractive possibility accounting for the observed enzymatic inhibition is that upon compound binding at the hydrophobic pocket, the conformation of FLN either becomes locked in a closed form or binding hampers the transition between closed and open forms of FLN.

### Genetic validations of FLN as an antimalaria drug target

Both biochemical and structural evidence suggest that FLN constitute a potential drug target. Given the FLN’s essentiality for the asexual growth of the parasite, we employed a conditional knock-down destabilization strategy to test this hypothesis. Hence, we generated a transgenic line with the *P. falciparum* 3D7 reference strain in which a chromosomal *fln* allele was fused with the HA tag and *Eschericia coli* dihydrofolate reductase destabilizing domain (DDD) (**Fig S4A, Fig S4B and S4C**). The *E. coli* DDD approach has been previously used to successfully disrupt gene function by protein destabilization in the absence of the DDD ligand trimethoprim (TMP) (Beck et al., 2014; Muralidharan et al., 2011; Muralidharan et al., 2012). By this approach, we achieved average reductions of the FLN protein levels by 83.6% from three biological replicates that was observed immediately after the TMP withdrawal. These reduced FLN levels were maintained for at least three subsequent intraerythrocytic developmental cycles (IDCs) with no effect on parasite growth or morphology (**Figs 4A and 4B, Fig. S4D** & **S4E**). Crucially, these partial depletions led to limited but significant increased susceptibilities of the transgenic parasites to the FLN interacting antimalarials including MMV000848, MK-4815, MFQ, and CQ (**Fig. 4C**). Specifically, we observed statistically significant decreases of IC50, ranging from ∼10% for MK-4815, MFQ, and CQ to ∼30% for MMV000848 (**Fig. 4C**). In contrast, the FLN knock down did not affect the *P. falciparum* susceptibilities to functionally unrelated antimalarial, dihydroartemisinin (**Fig. 4C**). As discussed below, the proteins destabilizations are currently the most suitable methods to study *P. falciparum* essential proteins; although it is often limited due to incomplete elimination of the target proteins from the cell (Ponpuak et al., 2007). This is indeed the case for FLN where we could reduce the protein level to ∼15% of its original levels. As such, the knock downs mediated only limited hypersensitivities. However, it is important to note that the achieved hypersensitivities were statistically significant for all four tested compounds for which we also observed their ability to bind and inhibit FLN.

**Figure 4.**
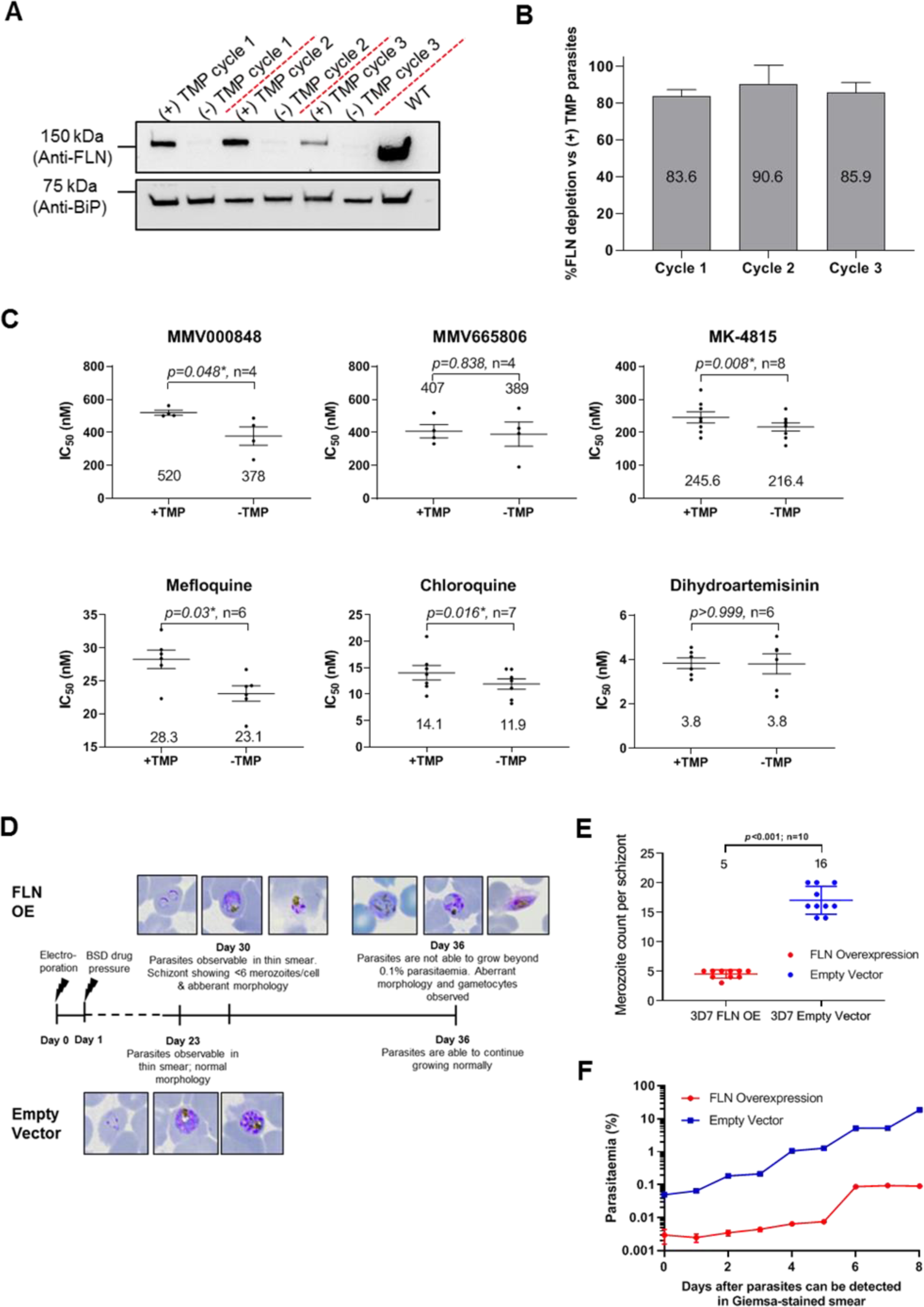
Characterization of FLN transfectant strains. **(A)** Western blot analysis of FLN levels in 3D7 FLN-DD in the presence or absence of TMP stabilizing ligand. Cycle number refers to the number of life cycle the parasites have been cultured after the initial removal of TMP. FLN were probed with anti-FLN antibody, whereas anti-PfBIP was used as loading control. **(B)** Densitometry analysis of FLN levels obtained from (A). Number in each bar in densitometry analysis represents average percentage decrease of FLN in parasites cultured without TMP stabilizing ligand compared to parasites cultured with TMP. Error bars represent standard deviation. Data are from 3 biological replicates. **(C)** Paired analysis of *in vitro* drug susceptibility of 3D7 FLN-DD against MMV000848, MMV665806, MK-4815, Mefloquine, CQ, and dihydroartemisinin in the presence or absence of TMP in the ring (top) or trophozoite (bottom) stage of the parasites. Numbers below the dots represents mean IC_50_. *P-values* were obtained from non-parametric Wilcoxon matched-pairs signed rank test, whereas *n* represent the number of biological replicates conducted for each experiment. * denotes statistically significant difference (i.e., *p<*0.05) between the pairs tested. (**D**) Timeline depicting the growth of FLN overexpression (FLN OE, top) and empty vector (control, bottom) strains after transfection. (**E)** Merozoite count from the schizonts of 3D7 FLN overexpression (blue) and empty vector (red) strains. Lines represent the mean of merozoite count per schizont (10 schizonts for each strain) and error bars represent standard deviation. Numbers represent median merozoite count in each strain. *P-*values were obtained from non-parametric Mann-Whitney test. **(F)** Measurement of parasitaemia following parasites recovery from initial drug pressure in FLN overexpression (red) or empty vector (blue) strains. Each point represents the mean value of measurements from two transfection attempts in Dd2 strain. Error bars which represent standard deviations are too small to be visible.

In addition to knock-down, we also investigated the effect of FLN overexpression by episomal overexpression of FLN. For that, we generated a transgenic cell line where full length FLN was fused to Hemagglutinin (HA) at the C-terminus and cloned into pBcamR_3xHA transfection vector providing adjustable expression via increased copy number induced by the selective marker responsive to blasticidin (Flueck et al., 2009). There, we examined the effect of FLN overexpression on drug sensitivities in a similar approach that we previously employed to study factors of artemisinin resistance (**Fig S4F**) (Rocamora et al., 2018). However, upon validation of overexpression (**Fig S4G and S4H)**, we observed a severe toxic effect caused by FLN overexpression represented by distorted parasite morphology of the trophozoite and schizont stages with ∼3-fold reductions in the number of newly formed merozoites (**Figs. 4D & E**). This is likely responsible for the severe growth impediment of the FLN overexpression line, keeping the overall parasitaemia below 0.1% (**Fig. 4F**). The FLN overexpression line also contained dramatically increased proportion of sexual stages (gametocytes) (**Fig. 4D**), a phenomenon commonly associated with stress (Josling et al., 2018). While the toxicity of FLN overexpression disallows our original plan of drug sensitivity measurements, it provides strong evidence of FLN’s critical role in *P. falciparum* asexual stage growth and development.

## Discussion

In this study we aimed to identify putative protein targets of quinolone-based drugs, historically the most successful class of antimalarials (Foley and Tilley, 1997). Despite decades of research, the MOAs of the quinoline antimalarials remain partly characterized with only few direct proteins targets fully verified (Dziekan et al., 2019; Read et al., 1999; Wong et al., 2017). The majority of quinoline drugs, including those studied here, are linked with the parasite DV where they exert their antiparasitic effect by interfering with the neutralizing polymerization of the toxic heme, released during hemoglobin digestion (hemozoin formation) (Foley and Tilley, 1997; Kapishnikov et al., 2021; Kapishnikov et al., 2019). This is particularly the case for CQ for which resistance is caused by lower accumulation in the DV to the levels that can no longer interfere with this process (reviewed in (Wicht et al., 2020)). Indeed, CQ and to a lesser degree AQ, LUM, and MFQ can induce high levels of free heme in the parasite cell likely as a part of their MOA (Combrinck et al., 2015; Combrinck et al., 2013; Herraiz et al., 2019). Nevertheless, there is mounting evidence that MOAs of quinoline involve additional mechanisms within the DV or other cellular compartments. For example, antimalarial activity of PYR was not only linked with hemozoin formation (Auparakkitanon et al., 2006) but also with inhibition of DNA topoisomerase II and possible other cellular processes (Chavalitshewinkoon et al., 1993) In our analysis PYR caused thermal shift of a set of sixty-six proteins that is highly enriched for factors of tRNA aminoacylation (**Fig. S5**). The biological role of tRNA aminoacylation is linked with several crucial pathways such as cytoplasmic translation, proteolysis, and amino acid metabolism all of which could be associated with the broad spectrum of the PYR MOAs (**Fig. S6**). In addition, several tRNA ligase/synthetases thermally stabilized by PYR appears to interact with other quinolines such as cysteine tRNA ligase (stabilized by CQ) (**Table S1**). At this stage, it is not clear whether the observed thermal stabilization was due to direct binding or downstream effect. However, the observed disruption in tRNA ligases could provide further clues to an extended view on quinoline MOAs that could be explored by future studies. Indeed, CQ been shown previously to induce dimerization of arginyl-tRNA ligase rendering the enzyme inactive (Jain et al., 2016). Moreover, tRNA ligases are being considered by several antimalarial development programs as high value drug targets (Herman et al., 2015; Hewitt et al., 2017; Istvan et al., 2023; Jain et al., 2017; Jain et al., 2015; Jain et al., 2016; Khan et al., 2013; Milne et al., 2022; Sharma et al., 2022; Sonoiki et al., 2016; Xie et al., 2022). Our MS-CETSA now links tRNA ligases with quinoline drug’s MOA suggesting their higher potential. In a similar fashion, the other putative targets of quinoline drug activities detected by our MS-CETSA could provide new avenues for further drug development.

In this study, we validated the interaction between FLN and five chemically diverse compounds. FLN is a M16C zinc metalloprotease whose main function is cleavage of small oligopeptides, typically products of upstream proteolytic processes (Ralph, 2007). FLN was originally defined as one of the proteases mediating hemoglobin degradation in the DV (Murata and Goldberg, 2003a, b). FLN is, however, distinct from the other hemoglobinases such as cysteine (falcipains), aspartate (plasmepsins) or histoaspartate (HAP) proteases (see below) (Chugh et al., 2013). FLN is by far the largest DV protease (130 kDa) exhibiting restricted and unique substrate peptide sequence specificities; preferring peptides with bulky charged residues (Eggleson et al., 1999b). Indeed, incapable to cleave hemoglobin or denatured globin, FLN cleaves 10-20 amino acid long peptides that are intermediate products of hemoglobin degradation (Klemba and Goldberg, 2002). This specialized role in hemoglobin digestion is a hallmark of FLN that carries inverted HXXEH motifs in its active site that is characteristic for M16 proteases in plants (Hooper, 1994; Rawlings and Barrett, 1995). Indeed, the closest phylogenetic ancestors of FLN are two Arabidopsis endopeptidase PreP1 and PreP2 that are responsible for degrading transit/targeting peptides in the chloroplast and mitochondria (Richter and Lamppa, 1998). FLN was found to also accumulate in the *Plasmodium* apicoplast, the endosymbiotic analog of chloroplast, and to a lesser degree in mitochondrion (Ponpuak et al., 2007). In the apicoplast, FLN cleaves free signal transit peptides of presumably >400 nuclear encoded proteins that are essential for its biochemical and cellular functions such as fatty acid type II synthesis, non-mevalonate isoprenoid synthesis, haem synthesis and iron-sulphur cluster assembly (McFadden and Yeh, 2017). While FLN accumulation in the plastid/mitochondria may represent its ancestral localization, its presence in the DV is possibly an evolutionary diversion of its role in trafficking process via endocytic pathway and as such may represent a functional diversification (Ralph, 2007). Nevertheless, in both locations, FLN functions as an endopeptidase, mediating organellar “clearance” of short polypeptides. As such, FLN inhibition can lead to massive accumulation of short polypeptides that could in turn disrupt the normal function of these organelles and consequently lead to parasite death. Moreover, inhibition of FLN in the DV can disrupt hemoglobin digestion, blocking the release of hemoglobin-derived amino acids that present a major metabolic source for *Plasmodium* parasite during its asexual development (Edgar et al., 2022).

Due to these unique aspects, FLN garnered some interest in the past few years as a promising antimalarial drug target (Eagon et al., 2021; Kahlon et al., 2021). Piperazine-derived hydroxamic acid inhibitors shown a particularly high promise as a new chemical scaffold targeting FLN in sub-micromolar concentrations (Chance et al., 2018; Kahlon et al., 2021). Subsequently, a virtual screen identified 34 new chemical structures with high probability of FLN binding, from which, eight have antimalarial activity (Eagon et al., 2021). All these current efforts, however, focused on the FLN proteolytic active site, in most cases relying on the drug’s ability to chelate the residing Zn^+2^. Here we identified an alternative inhibitory pocket that could potentially provide more effective disruption of its function(s) and not relying on a chelating effect. Indeed, targeting this newly discovered inhibitory pocket in FLN fulfils many criteria for drug target prioritization recently formulated by the Malaria Drug Accelerator (MalDA) consortium (Forte et al., 2021).

(i) *Chemical validations*. The MS-CETSA results indicated that the FLN-drug binding occurs in both lysate, indicating its chemical robustness, and *in-cell* suggesting its biological significance. This is consistent with our previous studies with other antimalarial drugs/inhibitors including pyrimethamine and E64, causing analogous thermal shifts in their putative targets PfDHFR and falcipain(s) respectively (Dziekan et al., 2019). Moreover, the biochemical and structural data provided here not only validated the observed FLN-drug binding but crucially, demonstrated its inhibition potential. From this perspective, FLN should be positioned into the antimalaria drug development pipeline along with several high priority candidates such as serine/threonine protein kinase (CLK3) (Alam et al., 2019), Acetyl CoA synthase (de Vries et al., 2022), tyrosine tRNA synthetase (Xie et al., 2022) farnesyl/geranylgeranyl pyrophosphate synthase (Gisselberg et al., 2018) and cGMP-dependent protein kinase (Baker et al., 2017). Like for all of these high priority targets, here we provide structural information of the inhibitor binding pocked that could guide further optimization using medicinal chemistry, including with the two MMV compounds.

*(ii.) Genetic Validation and Essentiality*; FLN is indispensable for *P. falciparum* asexual growth, which was previously demonstrated by a single gene approach (Ponpuak et al., 2007) but also in large scale deletion screens such as random piggyBac transposon mutagenesis in *P. falciparum* (Zhang et al., 2018) and screening of barcoded knockout *P. berghei* mutants (Bushell et al., 2017). Here, we showed that the conditional knock-down cannot eliminate all FLN from viable parasites but can reduce its level by >80%. This is consistent with previously studied essential genes for which the destabilization approaches typically fail to completely eliminate the protein, but could reduce their levels by up to ∼5-fold (Armstrong and Goldberg, 2007). For most essential proteins, even partial reductions are detrimental to parasite growth (Blomqvist et al., 2017; Dvorin et al., 2010). This was also demonstrated for a highly effective protein drug target, dihydrofolate reductase (PfDHFR) (Prommana et al., 2013) and targets of several experimental compounds that are at various stages of preclinical studies (Forte et al., 2021). Likewise, the destabilization knock-down of FLN lowered its levels down to ∼20-15% of its initial level representing a ∼5-fold reduction. However, even with these low FLN levels, *P. falciparum* parasites can progress through their IDC normally. Overexpression of FLN appears toxic to the parasite, which indicates a strict requirement of FLN expression levels for normal asexual development. This suggest that FLN essentiality correlates with certain expression levels for parasites growth in basal growing *in vitro* conditions. In the future it will be interesting to investigate its importance for parasite growth under varying stress conditions or natural infections. Overall, the partial conditional knock-down reported here did cause limited but statistically significant hypersensitivity of the *P. falciparum* asexual stages to four of the five verified FLN binders, including MK-4815, CQ, MFQ, and MMV000848. This is also consistent with the abovementioned drug discovery efforts in which conditional knock-down of the putative targets causes various levels of drug hypersensitivities indicating their direct involvements in their corresponding drug MOAs (Forte et al., 2021; Prommana et al., 2013).

*(ii.) Druggability* Here we show the ability of the FLN inhibitory pocket to bind several chemotypes which may indicate its binding “promiscuity”. In drug discovery, promiscuity is broadly defined as the ability of a single molecule to bind to multiple others (with low M affinities). From the small compounds perspective, promiscuity is referred to as PAINS (pan assay interfering compounds) that can to bind multiple protein targets and are therefore usually excluded from drug screening campaigns (Baell and Walters, 2014). Conversely, protein binding sites that are endowed with the capacity to bind several ligands (similar to FLN) provide high potential. One such example is represented by an esterase with a large binding cavity that can accept 60 structurally distinct substrates (Höppner et al., 2021) or the main protease of SARS-Cov2 (Ma et al., 2020). Similarly, FLN promiscuity chiefly results from the presence of a large substrate binding cavity that is fully accessible to the solvent in its open conformation. Hence, in addition to FLN natural peptide substrates, fused ring-bearing molecules with aliphatic moieties such as quinolines drugs or MMV000848 can easily diffuse and reach an exposed hydrophobic patch lined with Phe, Leu and Ile residues (**Fig. 3F**). We note that this type of promiscuity forms the basis for drug repurposing efforts. In this respect, the binding site appears as a druggable hotspot in a manner reminiscent to binding sites that can be uncovered using fragment-based screening. We also note the presence of several hydrophilic residues lining this otherwise hydrophobic binding site, such as Asp451 Asp526 and Asp509 or Glu530. These residues have the ability to form hydrogen bonds or salt bridges as is seen between Glu530, Asp451 and MK-4815 (**Fig. 3E**). Hence, these polar residues offer opportunities to provide polar interaction and impart specificity over the course of a medicinal chemistry program to improve inhibition capacity starting from the compounds studied here.

An important question raised by these studies is whether FLN is part of quinoline drug’s MOA, especially CQ? Our CETSA, ITC, and enzymatic assay data suggest that FLN inhibition by the quinolines CQ and MFQ require concentration at least three orders of magnitudes higher than its *in vitro* IC_50_ growth inhibition value. In the case of CQ, this is still highly compatible with FLN inhibition in the DV. There, CQ accumulates to sub-millimolar concentration, at least 1,000-fold more than that of the drug in the milieu (Famin and Ginsburg, 2002; Geary et al., 1986; Hawley et al., 1998). In addition several other observations link CQ’s MOA with FLN as a part of hemoglobin degradation (Eggleson et al., 1999a; Murata and Goldberg, 2003a, b). CQ was shown to induce a marked accumulation of undigested hemoglobin (Famin and Ginsburg, 2002), long-chain hemoglobin-derived peptides (Birrell et al., 2020) and di-/tri-peptides (Creek et al., 2016), indicating disruption in the hemoglobin digestion cascade. In the DV, FLN is likely to be present in a heme detoxification complex along with heme detoxification protein (HDP) and other proteases such as falcipains and plasmepsins (Chugh et al., 2013). The role of FLN in this complex is unknown. However, it is plausible that inhibition of FLN enzymatic activity might affect this HDP itself. Chugh et al. showed that hemoglobin degradation rate is higher than the rate of hemozoin crystal growth, suggesting a coupled feedback mechanism between these two processes to maintain non-toxic heme levels. Indeed, CQ can reduce hemoglobin binding to falcipain 2, which in turns blocks hemoglobin degradation and heme liberation (Chugh et al., 2013). It is possible that other proteases within HDP (such as FLN) regulate hemoglobin digestion and hemozoin formation (Goldberg, 2013). In this scenario, inhibition of FLN might lead to distortion of the HDP and with that disruption of the hemoglobin digestion/hemozoin regulatory feedback. Hence, it will be interesting to conduct biochemical study of FLN inhibition within the HDP context to decipher its overall regulatory role in heme processing.

## Materials and Methods

### Parasite culture

1. *P. falciparum* 3D7 cultures were maintained in Malaria Culture Media (MCM) containing RPMI-1640 medium (Gibco) supplemented with 0.25% Albumax II (Gibco), 0.1 mM hypoxanthine (Sigma), 0.2% sodium bicarbonate (Sigma), and 10 mg/L gentamycin. Human blood group O were used for parasite culture and cultures were maintained at 37°C with 5% CO_2_, 3% O_2_, and 92% N_2_. Culture media were replenished daily with freshly packed RBC added to the culture when necessary.

### Sample preparation for ITDR MS-CETSA and LC/MS analysis

Samples for intact-cell and lysate ITDR MS-CETSA and LC/MS analysis were prepared as described in Dziekan *et al* (Dziekan et al., 2020a; Dziekan et al., 2019). For both intact-cell and lysate MS-CETSA, synchronized mid-trophozoite stage (28±4 HPI) culture at 10% parasitaemia and 2% hematocrit was used. In case of intact-cell MS-CETSA, 1 mL of packed blood was loaded on MACS CS column (Milteny Biotech), washed, and eluted. Enriched infected RBC (iRBC)(>80% parasitaemia) were then incubated for 1 hour in MCM at 37°C with agitation for recovery and equilibration. The iRBC were then divided into 10 aliquots (75-90 million cells in 6 mL MCM for each aliquot) and treated with 4-fold dilution series of MMV000848, MMV665806, CQ, AQ, PYD, PIP, LUM at 10 µM top concentration, MK-4815 at 100 µM top concentration, or an equivalent volume of their respective solvent at 0.1% concentration, and incubated for 1 hour at 37°C with agitation. Cells were pelleted by centrifugation (2,500 rpm, 5 minutes), washed with PBS, resuspended in 150 µL PBS per aliquot and transferred in 50 µL aliquots to three 96-well plates, each representing different heat challenge temperature (37°C, 51°C, or 58°C). Samples were subjected to thermal challenge for 3 minutes, followed by 3 minutes cooling at 4°C. After heat treatment, 50 µL of lysis buffer (50 mM HEPES pH 7.5, 5 mM beta-glycerophosphate, 0.1 mM Na_3_VO_4_, 10 mM MgCl_2_, 2 mM TCEP, and cocktail EDTA-free protease inhibitors (Sigma)) were added to each well. Samples were then subjected to 3 times of freeze/thaw cycle, followed by mechanical shearing and centrifugation (20,000g, 20 minutes, 4°C) to isolate the soluble protein fraction. Protein quantification was then performed with bicinchonidic assay (BCA) protein assay kit (Pierce). In case of lysate MS-CETSA, the parasite culture was pelleted and incubated with 10x volume of 0.1% saponin in PBS (pH 7.2) for 5 minutes. Following centrifugation (2,500g, 5 minutes) the supernatant containing RBC cytosol was removed, whilst intact parasite pellet was washed 3 times with 50 mL of ice-cold PBS. Parasite was then resuspended in 1 mL of lysis buffer and subjected to 3 times of freeze/thaw cycle, followed by mechanical shearing. Parasite lysate was then centrifuged (20,000 g, 20 min, 4°C) and supernatant containing soluble parasite proteins was isolated. Protein concentration was quantified with BCA protein assay kit. Ten aliquots of 20 µg of proteins were added to MMV000848, MMV665806, AQ, PYD, PIP, LUM at 10 µM top concentration, MK-4815 at 250 µM top concentration, CQ at 500 µM top concentration, or an equivalent volume of their respective solvent at 0.1% concentration, incubated at room temperature for 3 minutes, and heated at 37°C, 51°C, or 57°C for 3 minutes followed by 3 minutes cooling at 4°C. The post heating lysates were centrifuged at 20,000 g for 20 minutes at 4°C and the supernatant was transferred to a fresh Eppendorf tube for further processing.

### Peptide preparation and labeling

Following quantification of lysate and intact-cell samples, the volume equivalent to 20 µg total protein in post heating treatment was aliquoted and incubated with denaturation and reduction buffer containing 100 mM triethylammonium bicarbonate (TEAB, pH 8.5), 20 mM tris(2-carboxyethyl)phosphine) (TCEP, pH 7.0), and 1% (w/v) Rapigest (Waters) at 55°C for 20 minutes, followed by alkylation with 55 mM chloroacetamide (CAA) at room temperature for 30 minutes. Samples were then sequentially digested with Lys-C (0.23 AU LysC/µg of protein, Wako) for 3 hours, followed by trypsin (0.05 µg trypsin/µg of protein) for 18 hours at 37°C. Enzyme activation and Rapigest degradation was performed by incubation with 1% trifluoroacetic acid (TFA, Sigma) for 45 minutes at 37°C and the sample supernatant containing peptides was collected by centrifugation at 20,000g for 10 minutes. The samples were then dried in a centrifugal vacuum evaporator and resolubilized with 200 mM TEAB to 1 µg/µl concentration. Peptides were labeled with TMT10plex Isobaric Label Reagent Set (Thermo Fisher Scientific) according to manufacturer’s protocol. Briefly, 4 µg of the digested protein was labeled for at least 1 hour at a condition of pH>6 and then quenched with 1 M Tris (pH 7.4). The labeled samples were subsequently pooled and desalted using a Oasis HLB column (Waters), followed by vacuum drying. Samples were resuspended in 5% (v/v) acetonitrile, 5% (v/v) ammonia and separated using high pH reverse phase Zorbax 300 extend C-18 4.6 mm x 250 mm (Agilent) column and liquid chromatography AKTA Micro system (GE Healthcare) was used for offline sample pre-fractionation into 96 fractions. The fractions were concatenated into 20 fractions and dried with a centrifugal vacuum concentrator.

### LC/MS analysis

The dried peptide sample fractions were resuspended in 1% acetonitrile, 0.5% (v/v) acetic acid, and 0.06% TFA in water prior to analysis on LC/MS. Online chromatography was performed using the reverse phase liquid chromatography Dionex 3000 UHPLC system coupled to a Q Exactive HF mass spectrometer (Thermo Fisher Scientific). Each fraction was separated on a 50 cm x 75 µm Easy-Spray analytical column (Thermo Fisher Scientific) in 70 minutes gradient of programmed mixture of mobile phase A (0.1% formic acid) and mobile phase B (99.9% acetonitrile, 0.1% formic acid) to the following gradient ober time: 1-55 min (2-25%), 55-57 min (25-50%), 57-58 min (50-85%), 58-63 min (85%), 63-70 min (2%). MS data were acquired using data-dependent acquisition (DDA) with full scan MS spectra acquired in the range of 350-1550 m/z at a resolution of 70,000 and AGC target of 3e6; Top12 MS^2^ 35,000 and AGC target of 1e5, and 1e5 isolation window at 1.2 m/z.

### Protein identification and quantification

Proteome Discoverer 2.1 software (Thermo Fisher Scientific) was used to perform protein identification and quantification based in the Xcalibur raw files. Raw spectra was submitted for database searching with Sequest HT (Thermo Scientific) search engine using combined Sanger Institute *P. falciparum* 3D7 or Dd2 Coding Sequences database and Uniprot database for human proteins. MS precursor mass tolerance was set at 30 ppm, fragment mass tolerance set at 0.06 Da and maximum missed cleavage sites of 3. Dynamic modifications include oxidation (M), deamidated (NQ), acetylation (N-terminal protein) while static modification includes carbamidomethyl (C). Forward/decoy searches were used for false discovery rate (FDR) estimation. Peptide and peptide spectrum matches (PSMs) with high=FDR 1% and medium=FDR 5% levels were accepted.

### Quantitative MS data analysis and MS-CETSA data processing

For downstream analysis, only unique peptides were used for quantification of associated protein abundance and only proteins quantified by at least 3 PSMs were used. Relative abundances of proteins under different compound concentrations were normalized against untreated control and fed into “mineCETSA” R package (available at https://github.com/nkdailingyun/mineCETSA) for data extraction, clean-up, normalization, curve fitting, hit selection, and plotting. The ITDR hit selection considers the global responses from the measured protein population and applies two thresholds: a Median Absolute Deviation scheme where the threshold was set at median ± 2.5*MAD and the R^2^ value of curve fitting (indicating goodness of fit) for the selection of the most prominent hits.

### Protein expression and purification

The Falcilysin gene (UniProtKB - Q76NL8, gene ID: 814283) coding for residues 59-1193 was synthesized and cloned into pNIC28-Bsa4 downstream a 6xHis tag and a TEV cleavage site (pNIC-FLN59-1193). BL21 (DE3) Rosetta T1R cells were transformed with pNIC-FLN59-1193 and grown at 37°C until OD_600nm_ reached 0.6. Protein expression was induced by addition of 0.5 mM IPTG for 16 hours at 16°C. The bacteria were harvested by centrifugation and cells were resuspended in 100 mM Na HEPES, 500 mM NaCl, 10 mM Imidazole, 10% (v/v) glycerol, 0.5 mM TCEP, pH 7.5, and lysed mechanically (LM20 microfluidizer). Lysate was cleared by centrifugation at 50,000 g for 1 h. Protein was purified by affinity chromatography using Ni-NTA beds (GE Healthcare) and eluted in 20 mM Na HEPES, 500 mM NaCl, 500 mM Imidazole, 10 % (v/v) glycerol, 0.5 mM TCEP, pH 7.5 before size exclusion chromatography using Hiload 16/600 Superdex 200 pg column (GE healthcare) equilibrated in 20 mM Na HEPES, 300 mM NaCl, 10% (v/v) glycerol, 0.5 mM TCEP, pH 7.5. Protein purity was checked by SDS-PAGE, before concentration by ultrafiltration to 22 mg.ml^-1^, flash frozen and stored at -80°C until use.

### Crystallization, X-ray Data Collection and Data Processing

Crystallization conditions for falcilysin were screened with a protein concentration of 22 mg⋅ml^-1^ using commercial kits (JCSG-plus^TM^ & Morpheus from Molecular Dimensions; Index & PEG/Ion Screen^TM^ from Hampton Research) using a mosquito crystallization robot (TTP Labtech). Crystals were obtained after a few days. To obtain the structures of protein-inhibitor complexes for CQ, mefloquine and MK-4815, native FLN crystals were soaked with each inhibitor at 10 mM concentration for approximately 1.5 h before being flash-frozen in liquid nitrogen. For MMV000848 and MMV665806, co-crystals were obtained by crystallization condition screening with a mixture of FLN (22 mg⋅ml^-1^) and compound (1 mM). Detailed buffer composition for each structure are presented in **Table S3**. X-ray diffraction intensities were collected at the PXIII and PXI beamlines (Swiss Light Source, Villigen), integrated and scaled using XDS (Kabsch, 2010). Structures were determined with molecular replacement as implemented in the program Phaser (McCoy et al., 2007) using the *P. falciparum* falcilysin ligand-free structure (Goldberg et al, unpublished, PDB access code: 3S5M) as a search probe. Refinement and simulated annealing omit map calculations were done using the Phenix package. Manual model corrections were performed at the computer graphics using Coot (Emsley and Cowtan, 2004). Data collection and refinement statistics are shown in **Table S3**. Structure similarity search was performed using the ssm server from EBI, (https://www.ebi.ac.uk/msd-srv/ssm/cgi-bin/ssmserver) (Krissinel and Henrick, 2004) and returned eight similar structures (with r.m.s.d. of 1.78-2.17 Å) all belonging to the M16C metalloprotease family. The volume of the catalytic cavity in the FLN protein was calculated using the 3V webserver (Voss and Gerstein, 2010). A scan of the Zn edge was performed to confirm the presence of Zinc ion in the active site of FLN crystals. A dataset was collected at 1.278 Å near the Zinc K edge and the anomalous signal was used to unambiguously confirm the presence and the location of the active-site Zinc ion.

### Isothermal titration microcalorimetry measurements

Isothermal titration calorimetry (ITC) data were collected using a MICROCAL PEAQ-ITC instrument (Malvern) at 25 °C with 19 injections and 500 rpm stirring speed in a buffer containing 20 mM Na-HEPES pH 7.5, 300 mM NaCl, 10% (*v/v*) glycerol, 0.5 mM TCEP, 1% DMSO. The K_d_ of MK-4815 (1.79 µM) was determined by direct titration of 200 µL of 100 µM FLN protein with 36.4 µL of 1 mM MK-4815 (**Fig 2A**). Determination of K_d_ of CQ to FLN via direct ITC proved challenging due to its low binding affinity and the limited sensitivity of ITC equipment. Instead, the K_d_ of CQ was estimated by a displacement titration assay (Sigurskjold, 2000), titrating a 200 µL mixture containing 100 µM FLN protein and 200 µM CQ with the injection of 36.4 µL of 1 mM MK-4815, which yielded an apparent K_d_ of 5.5 µM for MK-4815. The increase of apparent K_d_ for MK-4815 from 1.79 µM to 5.5 µM is caused by competition from CQ for the same binding site, from which a K_d_ of 116 µM for CQ was estimated using the competitive mode in Marven analysis software. Binding of MK-4815 to FLN was largely inhibited when a 200 µL mixture of 100 µM FLN protein and 2 mM CQ was titrated with injections of 36.4 µL of MK-4815 at 1 mM (**Fig S2A**). K_d_ values for MMV665806 and MMV000848 were measured by titrating 200 µL of 20 µM FLN protein with 36.4 µL of 200 µM respective MMV drugs. K_d_ of mefloquine was measured by titrating 200 µL of 100 µM FLN protein with 36.4 µL of 1 mM mefloquine. Negative control was performed by titrating 200 µL of 100 µM FLN protein with 36.4 µL of 1 mM artemether.

### Enzymatic assay

To determine IC_50_ values of the three inhibitors, FLN activity was monitored using substrate fluorescence quenching at pH 7.5 and pH 5.2 representing values encountered in the apicoplast/mitochondrion and DV respectively. The FLN protein at a concentration of 20 nM was incubated with 20 µM of substrate peptide (Murata and Goldberg, 2003b) (Dabcyl-YNHHS↓FFMEE-Edans where the arrow indicates the scissile peptide bond) in presence of various concentrations of inhibitors (ranging from 1mM to 0.48 µM for CQ and mefloquine and from 100 µM to 0.048 µM for MK-4815, from 500 µM to 244.14 nM for MMV665806 and MMV000848). Measurements for each concentration were performed as triplicate. A TECAN 10M plate reader, operated at 336 nm and 495 nm wavelengths for excitation and emission respectively was used to follow FLN protease activity. The reaction was started with automated addition of the enzyme into a mixture of substrate and inhibitor, followed for a total duration of 1 min. The initial velocity was determined automatically using Magellan® software provided by the manufacturer. The V_max_ values obtained were plotted against the log of the inhibitor concentration in GraphPad Prism to obtain the IC_50_ values. A negative control was performed with Lumefantrine at pH 7.5.

To assess the MOA of MK-4815, Michaelis-Menten steady state kinetics parameters were measured in the presence of various inhibitor concentrations. A serial dilution of substrate starting from a maximum value of 20 µM of substrate was incubated with three concentrations of inhibitor ranging from approximately 0.1 to 10 times of the K_d_ values (0.1 µM, 2 µM and 10 µM of MK-4815). The reaction was started with automated addition of the enzyme into a mixture comprising the substrate and inhibitor and followed for 1 min. The initial velocity was determined automatically using Magellan® software. The values of reciprocal velocity were plotted as a function of reciprocal concentration in a Lineweaver-Burk graph.

### Plasmid construction and transfection

To generate plasmid for overexpression study, full length FLN open reading frame (omitting the stop codon) was PCR amplified from *P. falciparum* 3D7 genomic DNA with primers 5’-TAAGCACCATGGATGAATTTAACAAAATTAAT-3’ and 5’-TGCTTAGCTAGCTTCTATTAATACCTTTTTA-3’ and cloned into the NcoI/NheI sites of pBcamR_3xHA in-frame with the coding sequences of 3xHA to generate the plasmid pFLN OE. To generate a chromosomal FLN-3HA-DD fusion, a 670 bp fragment at the 3’ end of the FLN open reading frame (omitting the stop codon) was PCR amplified from *P. falciparum* 3D7 genomic DNA with primers 5’-TAAGCACTCGAGTAAAGTAAATGATCCAAC-3’ and 5’-TGCTTACCTAGGTTCTATTAATACCTTTTTA-3’ and cloned into the XhoI/AvrII sites of pFLN-HADB in-frame with the coding sequences of 3xHA and *E. coli* destabilization domain. One hundred micrograms each of the resulting plasmid and phDHFR were co-electroporated into ring stage parasites of 3D7 parasites as previously described (Wu et al., 1995). Plasmid-containing parasites were selected with 5 µM TMP (Sigma) and 2.5 µg/mL Blasticidin (Sigma), followed by limiting dilution of surviving transfectants. Clones were screened by PCR (list of primers are listed in Table S4) and one clone with validated integration was selected for further analysis, designated 3D7-FLN DD.

### Western blotting

Western blots analysis was performed with parasite lysates generated from trophozoite of parasites cultured with or without TMP (**Fig. 4A**). For each well, 10 µg (for FLN-DD experiment) of protein lysates for each treatment condition was fractionated in 4-15% Mini-Protean TGX protein gels (Bio-Rad) and transferred to nitrocellulose membrane using Bio-Rad Trans-Blot Turbo Transfer System with “Turbo Mixed MW” settings (constant 2.5A for 7 minutes). Primary antibodies used are anti-FLN (rabbit) at 1:2000 and anti-PfBIP (rabbit) at 1:5000 in 5% nonfat milk in TBS-Tween (TBST). HRP-conjugated secondary antibodies (Biolegend) diluted at 1:2000 in 5% nonfat milk in TBST was used accordingly to their corresponding primary antibodies. Blot was developed with Clarity Western ECL system (Bio-Rad), and the image was generated and analyzed using LAS-3000 imager (Fujifilm)

### Immunofluorescence Assay

For immunofluorescence microscopy, cells were fixed with 4% formaldehyde and 0.00075% glutaraldehyde before quenching with 0.1 M glycine and washing with PBS. Permeabilization was done with 0.1% Triton X-100 and blocking was done in 3% bovine serum albumin. Primary and secondary antibodies were diluted in 3% BSA with the following dilutions: rat anti-HA (1:100), rabbit anti-FLN (1:200), anti-rat Alexa Fluor 647 (1:1000) and anti-rabbit Alexa Fluor 488 (1:1000). Primary antibody incubation was done for one hour at room temperature. Slides were subsequently washed three times with PBS before staining with secondary antibody at room temperature for 45 minutes. Slides were then air-dried and glass coverslips mounted with ProLong Gold Antifade with DAPI (Thermo Fisher Scientific). Samples were then imaged with the Zeiss Live Cell Observer II. Image processing and colocalization analysis was done in Zen Blue.

### *In vitro* Drug Susceptibility Assay

Prior to drug assays, 3D7-FLN DD parasites were divided into two groups: one grown in the presence of TMP and the other grown in the absence of TMP to allow depletion of FLN prior to drug assay. These parasites were allowed to grow for one life cycle (∼44 hours) prior to the start of the assay. Parasites at ∼10-16 HPI were dispensed to 96 well plates containing 12 serially diluted concentrations of MMV000848, MMV665806, MK-4815, CQ, and MFQ to make final 2% hematocrit and 0.8% parasitemia. The top concentration of compounds used in was 10,000 nM for MMV000848, MMV665806, and MK-4815, 1,000 nM for CQ and MFQ, and 50 nM for DHA followed by 2-fold serial dilution and a drug-free control. Parasites were incubated with the drug for 72 hours to allow all parasites to invade and proceed to the next cycle. The number of new, viable parasites in each well on the subsequent replication cycle was quantified by flow cytometry. Cells were stained using 50 μL of 8 μM Hoechst 33342 in PBS (pH 7.2) for 15 minutes at 37°C, followed by addition of ice-cold 200 μL of PBS. Cells were quantified using LSR Fortessa X-20 Flow Cytometer (BD Biosciences) using UV laser (355nm) and results were analyzed with FACS Diva Software (BD Biosciences). Dose-response curves and IC_50_ estimation were obtained using GraphPad Prism 8 four-parameter dose-response curve equation (GraphPad Software Inc.). All assays were performed in at least 3 biological triplicates (N stated in Fig. 4C) and additionally in technical duplicates per dose. Non-parametric t-test (paired) was used to calculate significance.

## Data availability

The data that support the findings of this study are available from the corresponding author upon request. Coordinates were deposited with the PDB. Accession codes for FLN-compound co-crystal structures are: 7DI7 (FLN-CQ), 7DIA (FLN-Mefloquine), 7DIJ (FLN-MK-4815), 8HO4 (FLN-MMV000848) and 8H05 (FLN-MMV665805).

## Acknowledgement

This work was supported by the Singapore Ministry of Education AcRF Tier 1 grant #RG34/19 (S); Tier 3 grant # MOE2019-T3-1-007 and Singapore National Medical Research Council grant #NMRC/OFIRG/0040/2017

## Author contributions

Z.B. and J.L. conceived and developed the idea for this article. G.W., J.Q.L., J.M.D., and A.E.S. wrote the manuscript and performed data analysis. G.W., J.Q.L., J.M.D., A.E.S., Z.C., N.E.B.Z., J.B., R.T.J.K., L.S.J., S.T., K.D.G., and H.Y. carried out experimental work for the experiments presented. G.W., J.Q.L., J.M.D., A.E.S., A.P., D.O., N.P., R.M.S., and P.N. contributed to the development of experimental protocol and reagents in this article. All authors provided feedback and changes to the manuscript. G.W., J.Q.L., J.M.D., J.L., and Z.B. wrote the manuscript.

## References

1. Alam, M.M., Sanchez-Azqueta, A., Janha, O., Flannery, E.L., Mahindra, A., Mapesa, K., Char, A.B., Sriranganadane, D., Brancucci, N.M.B., Antonova-Koch, Y., et al. (2019). Validation of the protein kinase PfCLK3 as a multistage cross-species malarial drug target. Science 365.

2. Allman, E.L., Painter, H.J., Samra, J., Carrasquilla, M., and Llinas, M. (2016). Metabolomic Profiling of the Malaria Box Reveals Antimalarial Target Pathways. Antimicrob Agents Chemother 60, 6635–6649.

3. Armstrong, C.M., and Goldberg, D.E. (2007). An FKBP destabilization domain modulates protein levels in Plasmodium falciparum. Nat Methods 4, 1007–1009.

4. Ashley, E.A., Dhorda, M., Fairhurst, R.M., Amaratunga, C., Lim, P., Suon, S., Sreng, S., Anderson, J.M., Mao, S., Sam, B., et al. (2014). Spread of artemisinin resistance in Plasmodium falciparum malaria. The New England journal of medicine 371, 411–423.

5. Auparakkitanon, S., Chapoomram, S., Kuaha, K., Chirachariyavej, T., and Wilairat, P. (2006). Targeting of hematin by the antimalarial pyronaridine. Antimicrob Agents Chemother 50, 2197–2200.

6. Baell, J., and Walters, M.A. (2014). Chemistry: Chemical con artists foil drug discovery. Nature 513, 481–483.

7. Baker, D.A., Stewart, L.B., Large, J.M., Bowyer, P.W., Ansell, K.H., Jiménez-Díaz, M.B., El Bakkouri, M., Birchall, K., Dechering, K.J., Bouloc, N.S., et al. (2017). A potent series targeting the malarial cGMP-dependent protein kinase clears infection and blocks transmission. Nature communications 8, 430.

8. Balikagala, B., Fukuda, N., Ikeda, M., Katuro, O.T., Tachibana, S.I., Yamauchi, M., Opio, W., Emoto, S., Anywar, D.A., Kimura, E., et al. (2021). Evidence of Artemisinin-Resistant Malaria in Africa. N Engl J Med 385, 1163–1171.

9. Baragana, B., Hallyburton, I., Lee, M.C., Norcross, N.R., Grimaldi, R., Otto, T.D., Proto, W.R., Blagborough, A.M., Meister, S., Wirjanata, G., et al. (2015). A novel multiple-stage antimalarial agent that inhibits protein synthesis. Nature 522, 315–320.

10. Beck, J.R., Muralidharan, V., Oksman, A., and Goldberg, D.E. (2014). PTEX component HSP101 mediates export of diverse malaria effectors into host erythrocytes. Nature 511, 592–595.

11. Birrell, G.W., Challis, M.P., De Paoli, A., Anderson, D., Devine, S.M., Heffernan, G.D., Jacobus, D.P., Edstein, M.D., Siddiqui, G., and Creek, D.J. (2020). Multi-omic Characterization of the Mode of Action of a Potent New Antimalarial Compound, JPC-3210, Against Plasmodium falciparum. Mol Cell Proteomics 19, 308–325.

12. Blomqvist, K., DiPetrillo, C., Streva, V.A., Pine, S., and Dvorin, J.D. (2017). Receptor for Activated C-Kinase 1 (PfRACK1) is required for Plasmodium falciparum intra-erythrocytic proliferation. Molecular and biochemical parasitology 211, 62–66.

13. Bushell, E., Gomes, A.R., Sanderson, T., Anar, B., Girling, G., Herd, C., Metcalf, T., Modrzynska, K., Schwach, F., Martin, R.E., et al. (2017). Functional Profiling of a Plasmodium Genome Reveals an Abundance of Essential Genes. Cell 170, 260–272.e268.

14. Chakrabarti, A., Narayana, C., Joshi, N., Garg, S., Garg, L.C., Ranganathan, A., Sagar, R., Pati, S., and Singh, S. (2022). Metalloprotease Gp63-Targeting Novel Glycoside Exhibits Potential Antileishmanial Activity. Frontiers in cellular and infection microbiology 12, 803048.

15. Challis, M.P., Devine, S.M., and Creek, D.J. (2022). Current and emerging target identification methods for novel antimalarials. International journal for parasitology Drugs and drug resistance 20, 135–144.

16. Chance, J.P., Fejzic, H., Hernandez, O., Istvan, E.S., Andaya, A., Maslov, N., Aispuro, R., Crisanto, T., Nguyen, H., Vidal, B., et al. (2018). Development of piperazine-based hydroxamic acid inhibitors against falcilysin, an essential malarial protease. Bioorganic & medicinal chemistry letters 28, 1846–1848.

17. Chavalitshewinkoon, P., Wilairat, P., Gamage, S., Denny, W., Figgitt, D., and Ralph, R. (1993). Structure-activity relationships and modes of action of 9-anilinoacridines against chloroquine-resistant Plasmodium falciparum in vitro. Antimicrob Agents Chemother 37, 403–406.

18. Chugh, M., Sundararaman, V., Kumar, S., Reddy, V.S., Siddiqui, W.A., Stuart, K.D., and Malhotra, P. (2013). Protein complex directs hemoglobin-to-hemozoin formation in Plasmodium falciparum. Proceedings of the National Academy of Sciences of the United States of America 110, 5392–5397.

19. Combrinck, J.M., Fong, K.Y., Gibhard, L., Smith, P.J., Wright, D.W., and Egan, T.J. (2015). Optimization of a multi-well colorimetric assay to determine haem species in Plasmodium falciparum in the presence of anti-malarials. Malar J 14, 253.

20. Combrinck, J.M., Mabotha, T.E., Ncokazi, K.K., Ambele, M.A., Taylor, D., Smith, P.J., Hoppe, H.C., and Egan, T.J. (2013). Insights into the role of heme in the mechanism of action of antimalarials. ACS Chem Biol 8, 133–137.

21. Cowell, A.N., Istvan, E.S., Lukens, A.K., Gomez-Lorenzo, M.G., Vanaerschot, M., Sakata-Kato, T., Flannery, E.L., Magistrado, P., Owen, E., Abraham, M., et al. (2018). Mapping the malaria parasite druggable genome by using in vitro evolution and chemogenomics. Science (New York, NY) 359, 191–199.

22. Creek, D.J., Chua, H.H., Cobbold, S.A., Nijagal, B., MacRae, J.I., Dickerman, B.K., Gilson, P.R., Ralph, S.A., and McConville, M.J. (2016). Metabolomics-Based Screening of the Malaria Box Reveals both Novel and Established Mechanisms of Action. Antimicrob Agents Chemother 60, 6650–6663.

23. de Vries, L.E., Jansen, P.A.M., Barcelo, C., Munro, J., Verhoef, J.M.J., Pasaje, C.F.A., Rubiano, K., Striepen, J., Abla, N., Berning, L., et al. (2022). Preclinical characterization and target validation of the antimalarial pantothenamide MMV693183. Nature communications 13, 2158.

24. Dondorp, A.M., Nosten, F., Yi, P., Das, D., Phyo, A.P., Tarning, J., Lwin, K.M., Ariey, F., Hanpithakpong, W., Lee, S.J., et al. (2009). Artemisinin resistance in Plasmodium falciparum malaria. N Engl J Med 361, 455–467.

25. Dvorin, J.D., Martyn, D.C., Patel, S.D., Grimley, J.S., Collins, C.R., Hopp, C.S., Bright, A.T., Westenberger, S., Winzeler, E., Blackman, M.J., et al. (2010). A plant-like kinase in Plasmodium falciparum regulates parasite egress from erythrocytes. Science (New York, NY) 328, 910–912.

26. Dziekan, J.M., Wirjanata, G., Dai, L., Go, K.D., Yu, H., Lim, Y.T., Chen, L., Wang, L.C., Puspita, B., Prabhu, N., et al. (2020a). Cellular thermal shift assay for the identification of drug-target interactions in the Plasmodium falciparum proteome. Nat Protoc.

27. Dziekan, J.M., Wirjanata, G., Dai, L., Go, K.D., Yu, H., Lim, Y.T., Chen, L., Wang, L.C., Puspita, B., Prabhu, N., et al. (2020b). Cellular thermal shift assay for the identification of drug-target interactions in the Plasmodium falciparum proteome. Nat Protoc 15, 1881–1921.

28. Dziekan, J.M., Yu, H., Chen, D., Dai, L., Wirjanata, G., Larsson, A., Prabhu, N., Sobota, R.M., Bozdech, Z., and Nordlund, P. (2019). Identifying purine nucleoside phosphorylase as the target of quinine using cellular thermal shift assay. Sci Transl Med 11.

29. Eagon, S., Howland, M., Heying, M., Callant, E., Brar, N., Pompa, E., and Mallari, J.P. (2021). Identification of Plasmodium falciparum falcilysin inhibitors by a virtual screen. Bioorg Med Chem Lett 52, 128394.

30. Edgar, R.C.S., Siddiqui, G., Hjerrild, K., Malcolm, T.R., Vinh, N.B., Webb, C.T., Holmes, C., MacRaild, C.A., Chernih, H.C., Suen, W.W., et al. (2022). Genetic and chemical validation of Plasmodium falciparum aminopeptidase PfA-M17 as a drug target in the hemoglobin digestion pathway. eLife 11.

31. Eggleson, K.K., Duffin, K.L., and Goldberg, D.E. (1999a). Identification and characterization of falcilysin, a metallopeptidase involved in hemoglobin catabolism within the malaria parasite Plasmodium falciparum. The Journal of biological chemistry 274, 32411–32417.

32. Eggleson, K.K., Duffin, K.L., and Goldberg, D.E. (1999b). Identification and characterization of falcilysin, a metallopeptidase involved in hemoglobin catabolism within the malaria parasite Plasmodium falciparum. The Journal of biological chemistry 274, 32411–32417.

33. Emsley, P., and Cowtan, K. (2004). Coot: model-building tools for molecular graphics. Acta crystallographica Section D, Biological crystallography 60, 2126–2132.

34. Famin, O., and Ginsburg, H. (2002). Differential effects of 4-aminoquinoline-containing antimalarial drugs on hemoglobin digestion in Plasmodium falciparum-infected erythrocytes. Biochem Pharmacol 63, 393–398.

35. Favuzza, P., de Lera Ruiz, M., Thompson, J.K., Triglia, T., Ngo, A., Steel, R.W.J., Vavrek, M., Christensen, J., Healer, J., Boyce, C., et al. (2020). Dual Plasmepsin-Targeting Antimalarial Agents Disrupt Multiple Stages of the Malaria Parasite Life Cycle. Cell host & microbe 27, 642–658.e612.

36. Flannery, E.L., McNamara, C.W., Kim, S.W., Kato, T.S., Li, F., Teng, C.H., Gagaring, K., Manary, M.J., Barboa, R., Meister, S., et al. (2015). Mutations in the P-type cation-transporter ATPase 4, PfATP4, mediate resistance to both aminopyrazole and spiroindolone antimalarials. ACS Chem Biol 10, 413–420.

37. Flueck, C., Bartfai, R., Volz, J., Niederwieser, I., Salcedo-Amaya, A.M., Alako, B.T., Ehlgen, F., Ralph, S.A., Cowman, A.F., Bozdech, Z., et al. (2009). Plasmodium falciparum heterochromatin protein 1 marks genomic loci linked to phenotypic variation of exported virulence factors. PLoS Pathog 5, e1000569.

38. Foley, M., and Tilley, L. (1997). Quinoline antimalarials: mechanisms of action and resistance. Int J Parasitol 27, 231–240.

39. Forte, B., Ottilie, S., Plater, A., Campo, B., Dechering, K.J., Gamo, F.J., Goldberg, D.E., Istvan, E.S., Lee, M., Lukens, A.K., et al. (2021). Prioritization of Molecular Targets for Antimalarial Drug Discovery. ACS infectious diseases 7, 2764–2776.

40. Gamo, F.J., Sanz, L.M., Vidal, J., de Cozar, C., Alvarez, E., Lavandera, J.L., Vanderwall, D.E., Green, D.V., Kumar, V., Hasan, S., et al. (2010). Thousands of chemical starting points for antimalarial lead identification. Nature 465, 305–310.

41. Geary, T.G., Jensen, J.B., and Ginsburg, H. (1986). Uptake of [3H]chloroquine by drug-sensitive and -resistant strains of the human malaria parasite Plasmodium falciparum. Biochem Pharmacol 35, 3805–3812.

42. Gisselberg, J.E., Herrera, Z., Orchard, L.M., Llinas, M., and Yeh, E. (2018). Specific Inhibition of the Bifunctional Farnesyl/Geranylgeranyl Diphosphate Synthase in Malaria Parasites via a New Small-Molecule Binding Site. Cell Chem Biol 25, 185–193 e185.

43. Goldberg, D.E. (2013). Complex nature of malaria parasite hemoglobin degradation [corrected]. Proc Natl Acad Sci U S A 110, 5283–5284.

44. Hart, E.M., Mitchell, A.M., Konovalova, A., Grabowicz, M., Sheng, J., Han, X., Rodriguez-Rivera, F.P., Schwaid, A.G., Malinverni, J.C., Balibar, C.J., et al. (2019). A small-molecule inhibitor of BamA impervious to efflux and the outer membrane permeability barrier. Proc Natl Acad Sci U S A 116, 21748–21757.

45. Hawley, S.R., Bray, P.G., Mungthin, M., Atkinson, J.D., O’Neill, P.M., and Ward, S.A. (1998). Relationship between antimalarial drug activity, accumulation, and inhibition of heme polymerization in Plasmodium falciparum in vitro. Antimicrob Agents Chemother 42, 682–686.

46. Herman, J.D., Pepper, L.R., Cortese, J.F., Estiu, G., Galinsky, K., Zuzarte-Luis, V., Derbyshire, E.R., Ribacke, U., Lukens, A.K., Santos, S.A., et al. (2015). The cytoplasmic prolyl-tRNA synthetase of the malaria parasite is a dual-stage target of febrifugine and its analogs. Sci Transl Med 7, 288ra277.

47. Herneisen, A.L., and Lourido, S. (2021). Thermal Proteome Profiling to Identify Protein-ligand Interactions in the Apicomplexan Parasite Toxoplasma gondii. Bio-protocol 11, e4207.

48. Herraiz, T., Guillén, H., González-Peña, D., and Arán, V.J. (2019). Antimalarial Quinoline Drugs Inhibit β-Hematin and Increase Free Hemin Catalyzing Peroxidative Reactions and Inhibition of Cysteine Proteases. Sci Rep 9, 15398.

49. Hewitt, S.N., Dranow, D.M., Horst, B.G., Abendroth, J.A., Forte, B., Hallyburton, I., Jansen, C., Baragaña, B., Choi, R., Rivas, K.L., et al. (2017). Biochemical and Structural Characterization of Selective Allosteric Inhibitors of the Plasmodium falciparum Drug Target, Prolyl-tRNA-synthetase. ACS infectious diseases 3, 34–44.

50. Hooper, N.M. (1994). Families of zinc metalloproteases. FEBS Lett 354, 1–6.

51. Höppner, A., Bollinger, A., Kobus, S., Thies, S., Coscolín, C., Ferrer, M., Jaeger, K.E., and Smits, S.H.J. (2021). Crystal structures of a novel family IV esterase in free and substrate-bound form. The FEBS journal 288, 3570–3584.

52. Hovlid, M.L., and Winzeler, E.A. (2016). Phenotypic Screens in Antimalarial Drug Discovery. Trends in parasitology 32, 697–707.

53. Ismail, H.M., Barton, V., Phanchana, M., Charoensutthivarakul, S., Wong, M.H., Hemingway, J., Biagini, G.A., O’Neill, P.M., and Ward, S.A. (2016). Artemisinin activity-based probes identify multiple molecular targets within the asexual stage of the malaria parasites Plasmodium falciparum 3D7. Proc Natl Acad Sci U S A 113, 2080–2085.

54. Istvan, E.S., Guerra, F., Abraham, M., Huang, K.S., Rocamora, F., Zhao, H., Xu, L., Pasaje, C., Kumpornsin, K., Luth, M.R., et al. (2023). Cytoplasmic isoleucyl tRNA synthetase as an attractive multistage antimalarial drug target. Sci Transl Med 15, eadc9249.

55. Jafari, R., Almqvist, H., Axelsson, H., Ignatushchenko, M., Lundback, T., Nordlund, P., and Martinez Molina, D. (2014). The cellular thermal shift assay for evaluating drug target interactions in cells. Nat Protoc 9, 2100–2122.

56. Jain, V., Yogavel, M., Kikuchi, H., Oshima, Y., Hariguchi, N., Matsumoto, M., Goel, P., Touquet, B., Jumani, R.S., Tacchini-Cottier, F., et al. (2017). Targeting Prolyl-tRNA Synthetase to Accelerate Drug Discovery against Malaria, Leishmaniasis, Toxoplasmosis, Cryptosporidiosis, and Coccidiosis. Structure 25, 1495–1505.e1496.

57. Jain, V., Yogavel, M., Oshima, Y., Kikuchi, H., Touquet, B., Hakimi, M.A., and Sharma, A. (2015). Structure of Prolyl-tRNA Synthetase-Halofuginone Complex Provides Basis for Development of Drugs against Malaria and Toxoplasmosis. Structure 23, 819–829.

58. Jain, V., Yogavel, M., and Sharma, A. (2016). Dimerization of Arginyl-tRNA Synthetase by Free Heme Drives Its Inactivation in Plasmodium falciparum. Structure 24, 1476–1487.

59. Josling, G.A., Williamson, K.C., and Llinás, M. (2018). Regulation of Sexual Commitment and Gametocytogenesis in Malaria Parasites. Annu Rev Microbiol 72, 501–519.

60. Kabsch, W. (2010). XDS. Acta crystallographica Section D, Biological crystallography 66, 125–132.

61. Kahlon, G., Lira, R., Masvlov, N., Pompa, E., Brar, N., Eagon, S., Anderson, M.O., Andaya, A., Chance, J.P., Fejzic, H., et al. (2021). Structure guided development of potent piperazine-derived hydroxamic acid inhibitors targeting falcilysin. Bioorg Med Chem Lett 32, 127683.

62. Kapishnikov, S., Hempelmann, E., Elbaum, M., Als-Nielsen, J., and Leiserowitz, L. (2021). Malaria Pigment Crystals: The Achilles’ Heel of the Malaria Parasite. ChemMedChem 16, 1515–1532.

63. Kapishnikov, S., Staalsø, T., Yang, Y., Lee, J., Pérez-Berná, A.J., Pereiro, E., Yang, Y., Werner, S., Guttmann, P., Leiserowitz, L., et al. (2019). Mode of action of quinoline antimalarial drugs in red blood cells infected by Plasmodium falciparum revealed in vivo. Proc Natl Acad Sci U S A 116, 22946–22952.

64. Khan, S., Garg, A., Camacho, N., Van Rooyen, J., Kumar Pole, A., Belrhali, H., Ribas de Pouplana, L., Sharma, V., and Sharma, A. (2013). Structural analysis of malaria-parasite lysyl-tRNA synthetase provides a platform for drug development. Acta Crystallogr D Biol Crystallogr 69, 785–795.

65. King, J.V., Liang, W.G., Scherpelz, K.P., Schilling, A.B., Meredith, S.C., and Tang, W.J. (2014). Molecular basis of substrate recognition and degradation by human presequence protease. Structure 22, 996–1007.

66. Klemba, M., and Goldberg, D.E. (2002). Biological roles of proteases in parasitic protozoa. Annu Rev Biochem 71, 275–305.

67. Krissinel, E., and Henrick, K. (2004). Secondary-structure matching (SSM), a new tool for fast protein structure alignment in three dimensions. Acta crystallographica Section D, Biological crystallography 60, 2256–2268.

68. Kuhen, K.L., Chatterjee, A.K., Rottmann, M., Gagaring, K., Borboa, R., Buenviaje, J., Chen, Z., Francek, C., Wu, T., Nagle, A., et al. (2014). KAF156 is an antimalarial clinical candidate with potential for use in prophylaxis, treatment, and prevention of disease transmission. Antimicrob Agents Chemother 58, 5060–5067.

69. Liang, W.G., Wijaya, J., Wei, H., Noble, A.J., Mancl, J.M., Mo, S., Lee, D., Lin King, J.V., Pan, M., Liu, C., et al. (2022). Structural basis for the mechanisms of human presequence protease conformational switch and substrate recognition. Nature communications 13, 1833.

70. Lu, K.-Y., Quan, B., Sylvester, K., Srivastava, T., Fitzgerald, M.C., and Derbyshire, E.R. (2020). *Plasmodium* chaperonin TRiC/CCT identified as a target of the antihistamine clemastine using parallel chemoproteomic strategy. Proceedings of the National Academy of Sciences 117, 5810–5817.

71. Luth, M.R., Gupta, P., Ottilie, S., and Winzeler, E.A. (2018). Using in Vitro Evolution and Whole Genome Analysis To Discover Next Generation Targets for Antimalarial Drug Discovery. ACS infectious diseases 4, 301–314.

72. Ma, C., Hu, Y., Townsend, J.A., Lagarias, P.I., Marty, M.T., Kolocouris, A., and Wang, J. (2020). Ebselen, Disulfiram, Carmofur, PX-12, Tideglusib, and Shikonin Are Nonspecific Promiscuous SARS-CoV-2 Main Protease Inhibitors. ACS pharmacology & translational science 3, 1265–1277.

73. Martinez Molina, D., Jafari, R., Ignatushchenko, M., Seki, T., Larsson, E.A., Dan, C., Sreekumar, L., Cao, Y., and Nordlund, P. (2013). Monitoring drug target engagement in cells and tissues using the cellular thermal shift assay. Science 341, 84–87.

74. Martinez Molina, D., and Nordlund, P. (2016). The Cellular Thermal Shift Assay: A Novel Biophysical Assay for In Situ Drug Target Engagement and Mechanistic Biomarker Studies. Annu Rev Pharmacol Toxicol 56, 141–161.

75. Matter, H., Nazaré, M., Güssregen, S., Will, D.W., Schreuder, H., Bauer, A., Urmann, M., Ritter, K., Wagner, M., and Wehner, V. (2009). Evidence for C-Cl/C-Br…pi interactions as an important contribution to protein-ligand binding affinity. Angew Chem Int Ed Engl 48, 2911–2916.

76. McCoy, A.J., Grosse-Kunstleve, R.W., Adams, P.D., Winn, M.D., Storoni, L.C., and Read, R.J. (2007). Phaser crystallographic software. Journal of applied crystallography 40, 658–674.

77. McFadden, G.I., and Yeh, E. (2017). The apicoplast: now you see it, now you don’t. International journal for parasitology 47, 137–144.

78. Milne, R., Wiedemar, N., Corpas-Lopez, V., Moynihan, E., Wall, R.J., Dawson, A., Robinson, D.A., Shepherd, S.M., Smith, R.J., Hallyburton, I., et al. (2022). Toolkit of Approaches To Support Target-Focused Drug Discovery for Plasmodium falciparum Lysyl tRNA Synthetase. ACS infectious diseases 8, 1962–1974.

79. Muralidharan, V., Oksman, A., Iwamoto, M., Wandless, T.J., and Goldberg, D.E. (2011). Asparagine repeat function in a Plasmodium falciparum protein assessed via a regulatable fluorescent affinity tag. Proc Natl Acad Sci U S A 108, 4411–4416.

80. Muralidharan, V., Oksman, A., Pal, P., Lindquist, S., and Goldberg, D.E. (2012). Plasmodium falciparum heat shock protein 110 stabilizes the asparagine repeat-rich parasite proteome during malarial fevers. Nature communications 3, 1310.

81. Murata, C.E., and Goldberg, D.E. (2003a). Plasmodium falciparum falcilysin: a metalloprotease with dual specificity. J Biol Chem 278, 38022–38028.

82. Murata, C.E., and Goldberg, D.E. (2003b). Plasmodium falciparum falcilysin: an unprocessed food vacuole enzyme. Molecular and biochemical parasitology 129, 123–126.

83. Murithi, J.M., Owen, E.S., Istvan, E.S., Lee, M.C.S., Ottilie, S., Chibale, K., Goldberg, D.E., Winzeler, E.A., Llinás, M., Fidock, D.A., et al. (2020). Combining Stage Specificity and Metabolomic Profiling to Advance Antimalarial Drug Discovery. Cell chemical biology 27, 158–171.e153.

84. Paquet, T., Le Manach, C., Cabrera, D.G., Younis, Y., Henrich, P.P., Abraham, T.S., Lee, M.C.S., Basak, R., Ghidelli-Disse, S., Lafuente-Monasterio, M.J., et al. (2017). Antimalarial efficacy of MMV390048, an inhibitor of Plasmodium phosphatidylinositol 4-kinase. Sci Transl Med 9.

85. Phyo, A.P., Ashley, E.A., Anderson, T.J.C., Bozdech, Z., Carrara, V.I., Sriprawat, K., Nair, S., White, M.M., Dziekan, J., Ling, C., et al. (2016). Declining Efficacy of Artemisinin Combination Therapy Against P. Falciparum Malaria on the Thai-Myanmar Border (2003–2013): The Role of Parasite Genetic Factors. Clin Infect Dis 63, 784–791.

86. Ponpuak, M., Klemba, M., Park, M., Gluzman, I.Y., Lamppa, G.K., and Goldberg, D.E. (2007). A role for falcilysin in transit peptide degradation in the Plasmodium falciparum apicoplast. Mol Microbiol 63, 314–334.

87. Powles, M.A., Allocco, J., Yeung, L., Nare, B., Liberator, P., and Schmatz, D. (2012). MK-4815, a potential new oral agent for treatment of malaria. Antimicrob Agents Chemother 56, 2414–2419.

88. Prommana, P., Uthaipibull, C., Wongsombat, C., Kamchonwongpaisan, S., Yuthavong, Y., Knuepfer, E., Holder, A.A., and Shaw, P.J. (2013). Inducible knockdown of Plasmodium gene expression using the glmS ribozyme. PLoS One 8, e73783.

89. Ralph, S.A. (2007). Subcellular multitasking - multiple destinations and roles for the Plasmodium falcilysin protease. Molecular microbiology 63, 309–313.

90. Rawlings, N.D., and Barrett, A.J. (1995). Evolutionary families of metallopeptidases. Methods Enzymol 248, 183–228.

91. Read, J.A., Wilkinson, K.W., Tranter, R., Sessions, R.B., and Brady, R.L. (1999). Chloroquine binds in the cofactor binding site of Plasmodium falciparum lactate dehydrogenase. J Biol Chem 274, 10213–10218.

92. Richter, S., and Lamppa, G.K. (1998). A chloroplast processing enzyme functions as the general stromal processing peptidase. Proceedings of the National Academy of Sciences of the United States of America 95, 7463–7468.

93. Rocamora, F., Zhu, L., Liong, K.Y., Dondorp, A., Miotto, O., Mok, S., and Bozdech, Z. (2018). Oxidative stress and protein damage responses mediate artemisinin resistance in malaria parasites. PLoS Pathog 14, e1006930.

94. Rottmann, M., McNamara, C., Yeung, B.K., Lee, M.C., Zou, B., Russell, B., Seitz, P., Plouffe, D.M., Dharia, N.V., Tan, J., et al. (2010). Spiroindolones, a potent compound class for the treatment of malaria. Science (New York, NY) 329, 1175–1180.

95. Sharma, M., Mutharasappan, N., Manickam, Y., Harlos, K., Melillo, B., Comer, E., Tabassum, H., Parvez, S., Schreiber, S.L., and Sharma, A. (2022). Inhibition of Plasmodium falciparum phenylalanine tRNA synthetase provides opportunity for antimalarial drug development. Structure 30, 962–972.e963.

96. Sigurskjold, B.W. (2000). Exact analysis of competition ligand binding by displacement isothermal titration calorimetry. Analytical biochemistry 277, 260–266.

97. Sonoiki, E., Palencia, A., Guo, D., Ahyong, V., Dong, C., Li, X., Hernandez, V.S., Zhang, Y.K., Choi, W., Gut, J., et al. (2016). Antimalarial Benzoxaboroles Target Plasmodium falciparum Leucyl-tRNA Synthetase. Antimicrob Agents Chemother 60, 4886–4895.

98. Spangenberg, T., Burrows, J.N., Kowalczyk, P., McDonald, S., Wells, T.N., and Willis, P. (2013). The open access malaria box: a drug discovery catalyst for neglected diseases. PLoS One 8, e62906.

99. Spillman, N.J., Allen, R.J., McNamara, C.W., Yeung, B.K., Winzeler, E.A., Diagana, T.T., and Kirk, K. (2013). Na(+) regulation in the malaria parasite Plasmodium falciparum involves the cation ATPase PfATP4 and is a target of the spiroindolone antimalarials. Cell Host Microbe 13, 227–237.

100. Su, X.Z., and Miller, L.H. (2015). The discovery of artemisinin and the Nobel Prize in Physiology or Medicine. Science China Life sciences 58, 1175–1179.

101. Summers, R.L., Pasaje, C.F.A., Pisco, J.P., Striepen, J., Luth, M.R., Kumpornsin, K., Carpenter, E.F., Munro, J.T., Lin, D., Plater, A., et al. (2022). Chemogenomics identifies acetyl-coenzyme A synthetase as a target for malaria treatment and prevention. Cell chemical biology 29, 191–201.e198.

102. van der Pluijm, R.W., Imwong, M., Chau, N.H., Hoa, N.T., Thuy-Nhien, N.T., Thanh, N.V., Jittamala, P., Hanboonkunupakarn, B., Chutasmit, K., Saelow, C., et al. (2019). Determinants of dihydroartemisinin-piperaquine treatment failure in Plasmodium falciparum malaria in Cambodia, Thailand, and Vietnam: a prospective clinical, pharmacological, and genetic study. Lancet Infect Dis 19, 952–961.

103. van Schalkwyk, D.A. History of Antimalarial Agents. In eLS, pp. 1–5.

104. Voss, N.R., and Gerstein, M. (2010). 3V: cavity, channel and cleft volume calculator and extractor. Nucleic acids research 38, W555–562.

105. Wang, J., Zhang, C.J., Chia, W.N., Loh, C.C., Li, Z., Lee, Y.M., He, Y., Yuan, L.X., Lim, T.K., Liu, M., et al. (2015). Haem-activated promiscuous targeting of artemisinin in Plasmodium falciparum. Nature communications 6, 10111.

106. Wicht, K.J., Mok, S., and Fidock, D.A. (2020). Molecular Mechanisms of Drug Resistance in Plasmodium falciparum Malaria. Annu Rev Microbiol 74, 431–454.

107. Wong, W., Bai, X.C., Sleebs, B.E., Triglia, T., Brown, A., Thompson, J.K., Jackson, K.E., Hanssen, E., Marapana, D.S., Fernandez, I.S., et al. (2017). Mefloquine targets the Plasmodium falciparum 80S ribosome to inhibit protein synthesis. Nature microbiology 2, 17031.

108. World Health Organization (2021). World Malaria Report 2021 (Geneva, Switzerland).

109. Wu, Y., Sifri, C.D., Lei, H.H., Su, X.Z., and Wellems, T.E. (1995). Transfection of Plasmodium falciparum within human red blood cells. Proc Natl Acad Sci U S A 92, 973–977.

110. Xie, S.C., Gillett, D.L., Spillman, N.J., Tsu, C., Luth, M.R., Ottilie, S., Duffy, S., Gould, A.E., Hales, P., Seager, B.A., et al. (2018). Target Validation and Identification of Novel Boronate Inhibitors of the Plasmodium falciparum Proteasome. J Med Chem 61, 10053–10066.

111. Xie, S.C., Metcalfe, R.D., Dunn, E., Morton, C.J., Huang, S.C., Puhalovich, T., Du, Y., Wittlin, S., Nie, S., Luth, M.R., et al. (2022). Reaction hijacking of tyrosine tRNA synthetase as a new whole-of-life-cycle antimalarial strategy. Science 376, 1074–1079.

112. Yang, T., Ottilie, S., Istvan, E.S., Godinez-Macias, K.P., Lukens, A.K., Baragaña, B., Campo, B., Walpole, C., Niles, J.C., Chibale, K., et al. (2021). MalDA, Accelerating Malaria Drug Discovery. Trends in parasitology 37, 493–507.

113. Zhang, M., Wang, C., Otto, T.D., Oberstaller, J., Liao, X., Adapa, S.R., Udenze, K., Bronner, I.F., Casandra, D., Mayho, M., et al. (2018). Uncovering the essential genes of the human malaria parasite Plasmodium falciparum by saturation mutagenesis. Science 360.

