## Supplementary figures and tables for "Identification of an inhibitory pocket in falcilysin provides a new avenue for malaria drug development"

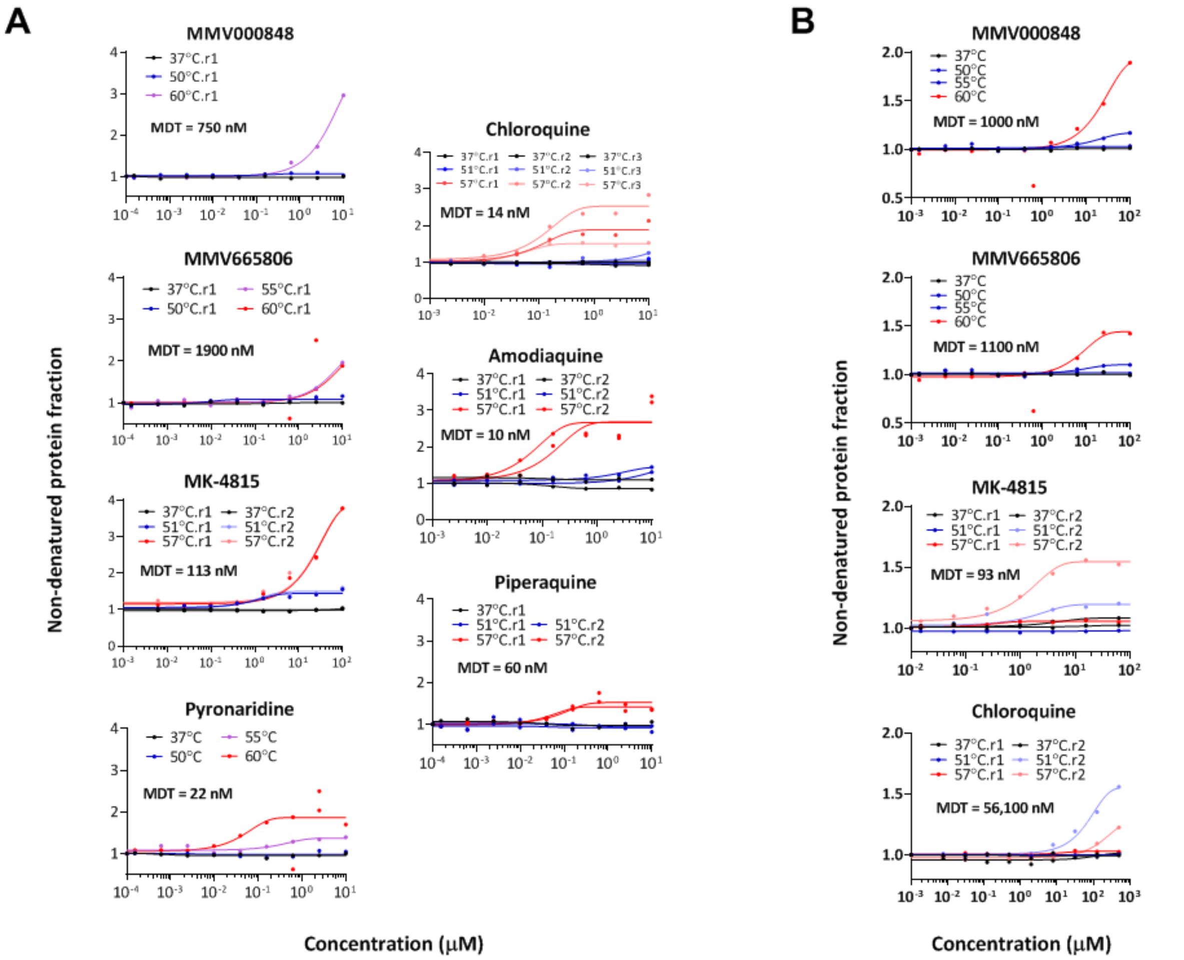


**Figure S1. Thermal stabilization profile of FLN**. Stabilization under thermal challenges (coloured) is plotted relative to no-drug control with non-denaturing control (37°C) plotted in black. Minimal dose threshold (MDT) stated are from ITDR 57°C (for MK-4815, chloroquine, amodiaquine, and piperaquine) or ITDR 60°C (MMV000848, MMV665806, and pyronaridine). Stabilization under thermal challenges (coloured) is plotted relative to no-drug control with non-denaturing control (37°C) plotted in black.

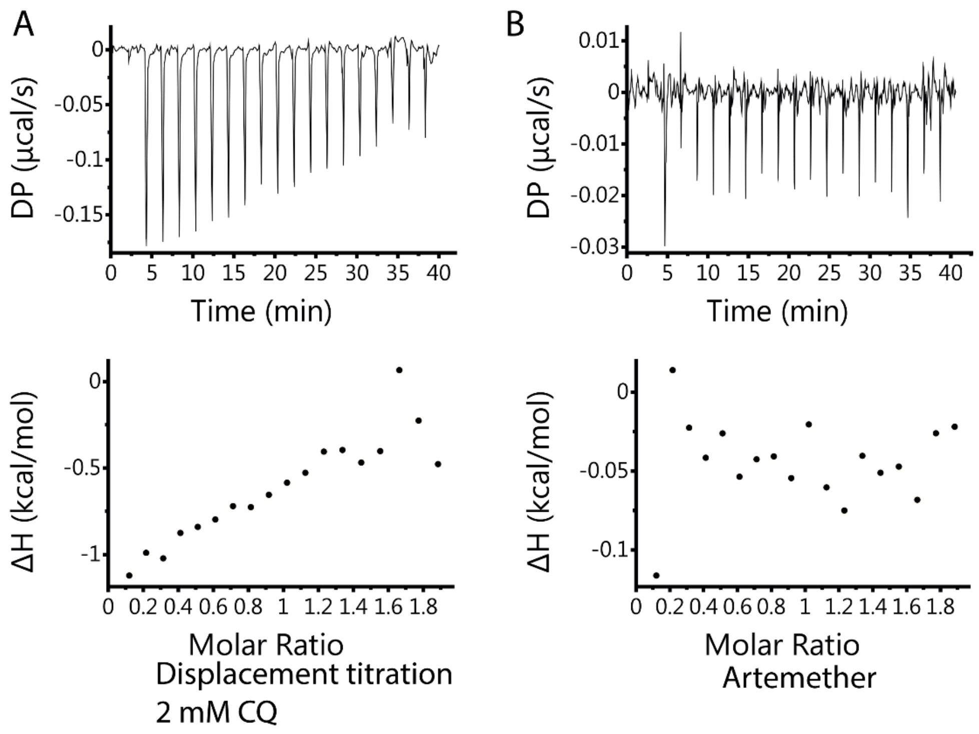


**Figure S2. Isothermal titration calorimetry measurements between FLN and CQ and artemether.** **(A)** Displacement titration of a mixture of 100 µM FLN and 2 mM of CQ with injection of 1 mM MK-4815. Due to the high concentration of CQ in the pre-mixture with FLN, the signal corresponding to MK-4815 binding upon injection was greatly reduced compared to the signal in the absence of CQ in the pre-mix. **(B)** Titration of 100 µM FLN with 1 mM artemether. No signal corresponding to the binding of artemether was detected, suggesting some binding specificity for FLN.

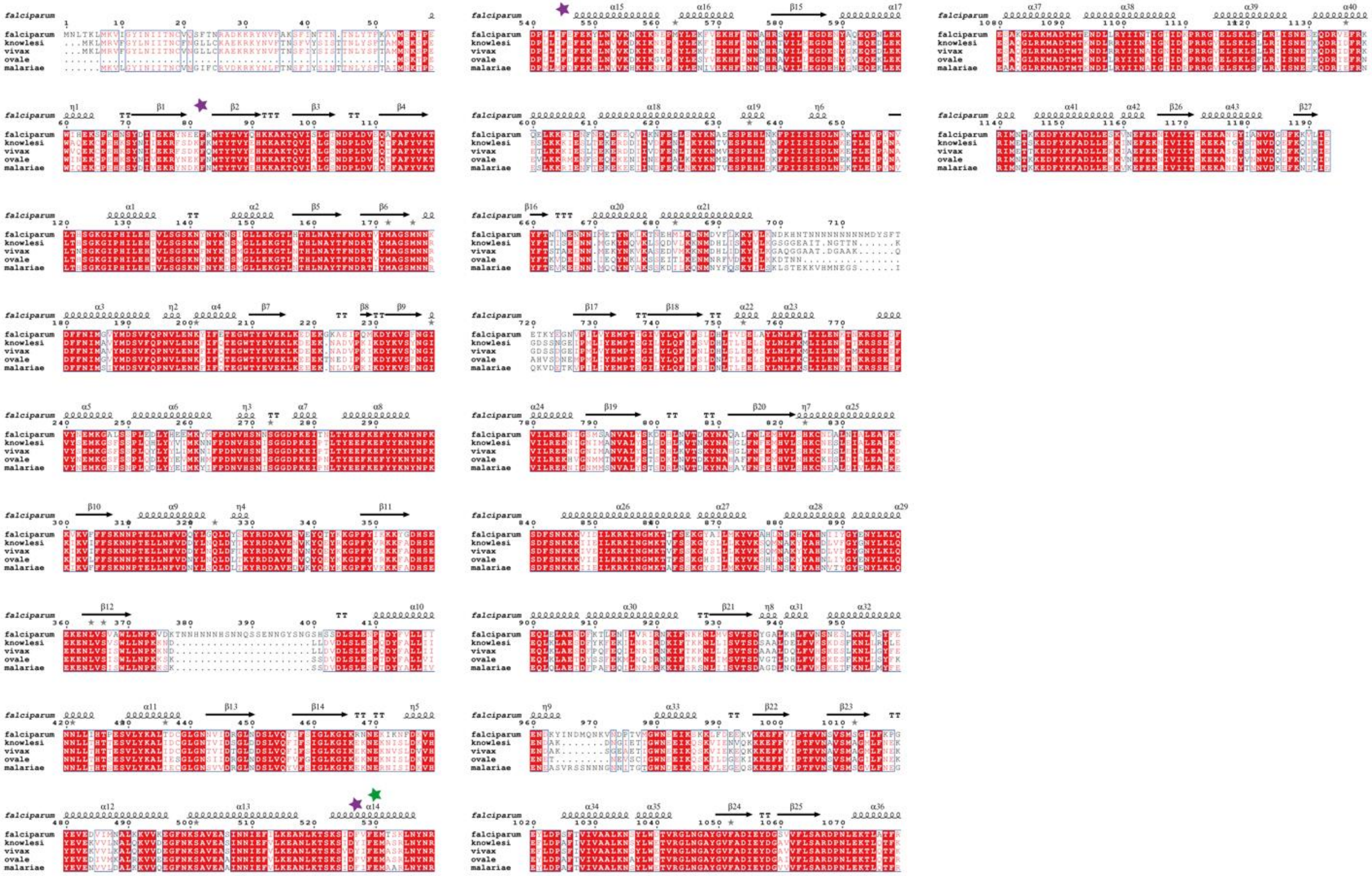

**Figure S3**. **Comparison of FLN sequence across different *Plasmodium* species**. Sequences of FLN genes from P. *falciparum* (PF3D7_1360800), P. *ovale* (PowCR01_110013700), P. *knowlesii* (A0A1Y3DII7|), P. *malariae* (A0A1D3SNQ8|) and P. *vivax* (A5K2L4|A5K2L4_PLAVS) were aligned. F82, F527 and F545 are featured with purple stars; E530 is featured with a green star. Secondary structure information is mapped based on *P. falciparum* FLN 3D crystal structure (this work). The structural alignment was made using ESPript3 server [1] at http://espript.ibcp.fr/ESPript/ESPript/.

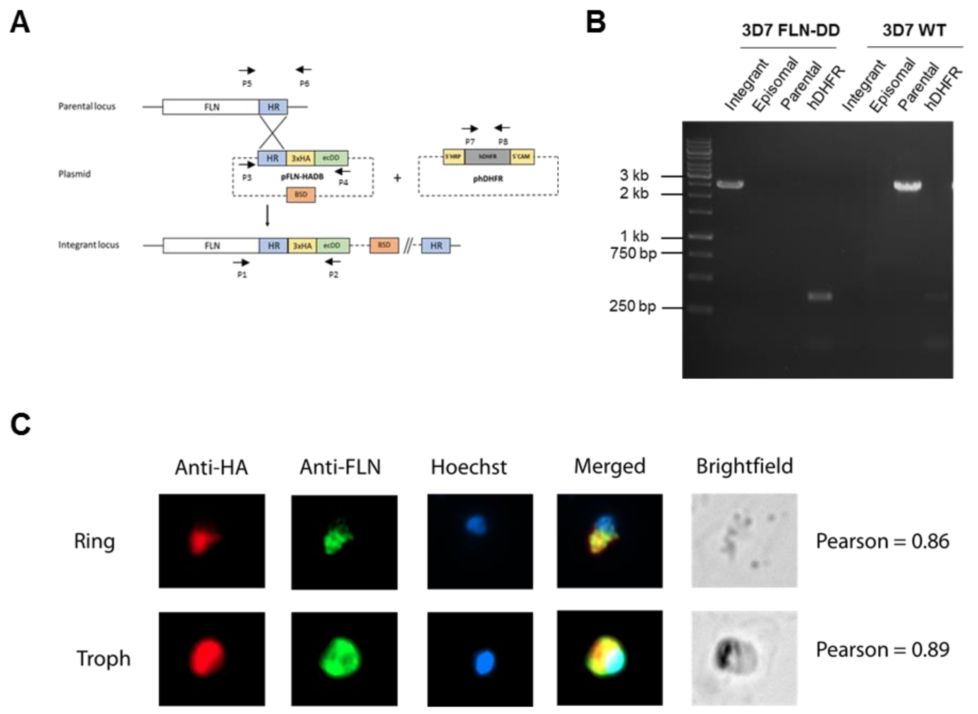


D

| Cycle no. | Multiplication Rate[a] | |
|---|---|---|
| | + TMP | - TMP |
| 1 | 5.2 (1.0) | 5.0 (0.5) |
| 2 | 5.2 (0.9) | 5.1 (1.0) |
| 3 | 5.1 (0.8) | 5.3 (0.5) |

[a]Numbers represent average parasite multiplication rate, whereas numbers in brackets represent standard deviation. Data are from 4 biological replicates

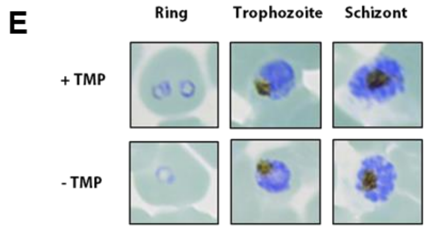


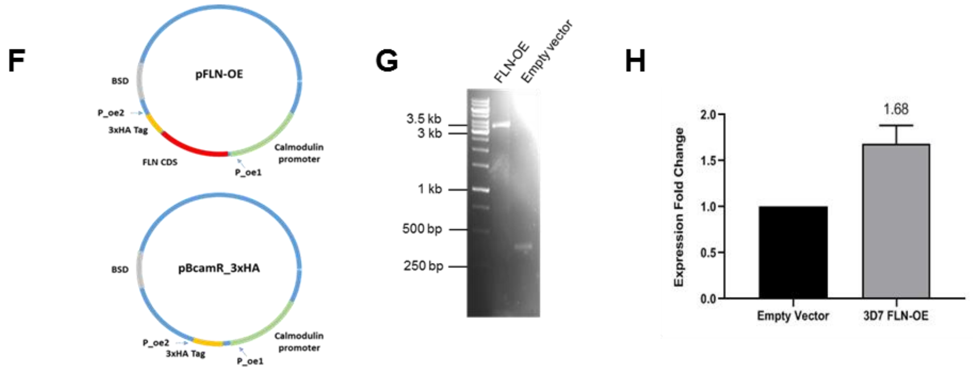


**Figure S4**. **Transfection strategy and validation of transgenic FLN line**. **(A)** 3D7 wild type strain was transfected with two constructs: 1) pFLN-HADB containing FLN homology region, 3x hemagglutinin (HA) tag, and *E.coli* destabilization domain (DD) and 2) phDHFR encoding for human dihydrofolate reductase (hDHFR) to confer resistance to trimethoprim (TMP). pFLN-HADB construct was integrated to parental locus via single site crossover, whereas phDHFR was retained as episomal plasmid and hDHFR was expressed with calmodulin promoter. p1-p8 denotes primer location for PCR validation. **(B)** PCR validation of 3D7 FLN-DD line. Integrant, episomal, parental, and hDHFR specific product was amplified using primer pair p1/p2, p3/4. p5/p6, and p7/p8, respectively. Primer sequences are listed in Table Sx. **(C)** Immunofluorescence assay (IFA) to check proper integration of FLN-HA. Ring (top) and trophozoite (bottom) stages were probed with anti-HA (red) and anti-FLN (green). Pearson value was obtained by plotting the signals from anti-HA and anti FLN. **(D)** Multiplication rate of 3D7 FLN-DD cultured with or without TMP. **(E)** Giemsa-stained micrograph of 3D7 FLN-DD cultured with or without TMP. **(F)** Plasmid design for FLN episomal overexpression study. pBcamR 3xHA was used to generate“empty control” line, whereas pFLN-OE was used to generate “FLN overexpression” line. **(G)** PCR validation of FLN-OE line. Episomal FLN specific product was amplified using primer pair p_oe1/p_oe2. Primer sequences are listed in table Sx. **(H)** qPCR gene expression analysis of FLN OE line. Y-axis denotes expression fold-change of FLN-OE strain compared to empty vector. Number denotes average and error bars represent standard deviation of fold expression changes across 3 independent replicates.

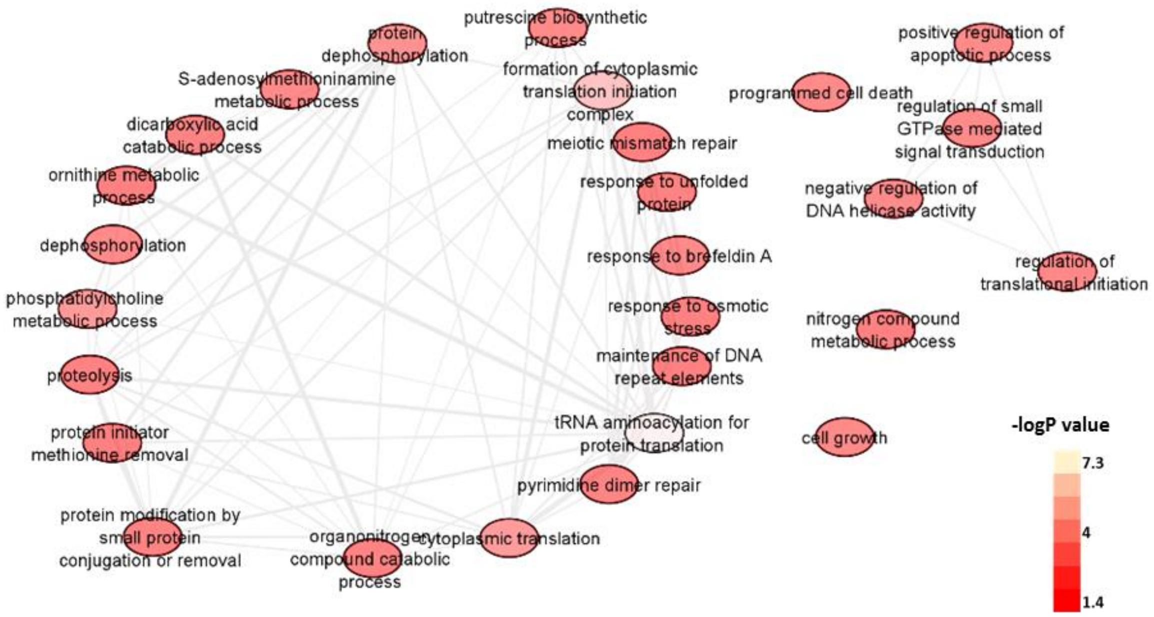


**Figure S5**. **Pathway over-representation analysis of proteins identified as hits in in-cell PYD-treated parasites**. Enriched pathways from protein datasets were obtained from PlasmoDB pathway enrichment tool based on Gene Ontology (GO) biological processes database with *p*-value cutoff set at 0.05. Nodes and edges were based on computed and curated pathways.

### Table S1. Proteins Identified as Protein Hits in Intact Cell CETSA

| Drug | Gene ID | Description | Heat challenge (°C) | MDT (nM) | *In vitro* $IC_{50}$ (nM) | MDT/$IC_{50}$ ratio | Median MDT/$IC_{50}$ ratio |
|---|---|---|---|---|---|---|---|
| **Amodiaquine 51°C; N=681 proteins 57°C; N=578 proteins** | PF3D7_0320900 | histone H2A.Z | 57 | 3610.84 | 35.00 | 103.17 | 2.96 |
| | PF3D7_0528200 | eukaryotic translation initiation factor 3 subunit E putative | 57 | 54.78 | | 1.57 | |
| | PF3D7_0624000 | hexokinase | 57 | 152.18 | | 4.35 | |
| | PF3D7_1360800 | falcilysin | 57 | 9.25 | | 0.26 | |
| **Chloroquine 51°C; N=802 proteins 57°C; N=694 proteins** | PF3D7_0708400 | heat shock protein 90 | 51 | 24.49 | 25.00 | 0.98 | 0.82 |
| | PF3D7_1015200 | cysteine--tRNA ligase | 57 | 17.74 | | 0.71 | |
| | PF3D7_1130400 | 26S protease regulatory subunit 6A putative | 57 | 0.78 | | 0.03 | |
| | PF3D7_1132000 | ubiquitin-like protein putative | 51 | 147.68 | | 5.91 | |
| | PF3D7_1233600 | asparagine and aspartate rich protein 1 | 51 | 12.18 | | 0.49 | |
| | PF3D7_1359400 | CUGBP Elav-like family member 1 | 51 | 24.31 | | 0.97 | |
| | PF3D7_1360800 | falcilysin | 57 | 13.57 | | 0.54 | |
| | PF3D7_1423200 | peptidyl-prolyl cis-trans isomerase | 51 | 38.09 | | 1.52 | |
| | PF3D7_1434300 | Hsp70/Hsp90 organizing protein | 51 | 20.55 | | 0.82 | |
| **Lumenfantrine 50°C; N=661 proteins 55°C; N=573 proteins 60°C; N=507 proteins** | PF3D7_1135400 | hotdog-domain containing protein | 60 | 410.34 | 20.00 | 20.52 | 20.52 |
| **MK-4815 51°C; N=616 proteins 57°C; N=655 proteins** | PF3D7_0305600 | DNA-(apurinic or apyrimidinic site) lyase putative | 57 | 12828.30 | 400.00 | 32.07 | 2.59 |
| | PF3D7_0708400 | heat shock protein 90 | 57 | 460.73 | | 1.15 | |
| | PF3D7_0814000 | 60S ribosomal protein L13-2 putative | 57 | 1546.14 | | 3.87 | |
| | PF3D7_1124700 | GrpE protein homolog mitochondrial putative | 57 | 6807.58 | | 17.02 | |
| | PF3D7_1212700 | eukaryotic translation initiation factor 3 subunit A putative | 51 | 8051.39 | | 20.13 | |
| | PF3D7_1222300 | endoplasmin putative | 57 | 1067.82 | | 2.67 | |
| | PF3D7_1360800 | falcilysin | 51 | 45.77 | | 0.11 | |
| | PF3D7_1360800 | falcilysin | 57 | 113.04 | | 0.28 | |
| | PF3D7_1401800 | choline kinase | 57 | 31.15 | | 0.08 | |
| | PF3D7_1434800 | mitochondrial acidic protein MAM33 putative | 57 | 1000.45 | | 2.50 | |
| **MMV000848 50°C; N=1166 proteins 60°C; N=825 proteins** | PF3D7_0110800 | transcription initiation factor TFIIB putative | 60 | 196.53 | 500.00 | 0.39 | 1.24 |
| | PF3D7_0204700 | hexose transporter | 50 | 316.49 | | 0.63 | |
| | PF3D7_0311300 | phosphatidylinositol 3- and 4-kinase putative | 60 | 985.85 | | 1.97 | |
| | PF3D7_0312800 | 60S ribosomal protein L26 putative | 50 | 189.79 | | 0.38 | |
| | PF3D7_0407800 | conserved Plasmodium protein unknown function | 60 | 609.72 | | 1.22 | |
| | PF3D7_0409400 | chaperone protein DnaJ | 60 | 1605.60 | | 3.21 | |
| | PF3D7_0505500 | DNA mismatch repair protein MSH6 putative | 60 | 383.36 | | 0.77 | |
| | PF3D7_0505700 | conserved Plasmodium membrane protein unknown function | 60 | 3846.84 | | 7.69 | |
| | PF3D7_0513300 | purine nucleoside phosphorylase | 60 | 22.23 | | 0.04 | |
| | PF3D7_0526200 | ADP-ribosylation factor GTPase-activating protein putative | 50 | 1290.89 | | 2.58 | |

| | | | | | | | |
|---|---|---|---|---|---|---|---|
| | PF3D7_0611600 | basal complex transmembrane protein 1 | 60 | 351.65 | | 0.70 | |
| | PF3D7_0710600 | 60S ribosomal protein L34 | 50 | 339.18 | | 0.68 | |
| | PF3D7_0819600 | conserved Plasmodium protein unknown function | 50 | 1440.99 | | 2.88 | |
| | PF3D7_0822900 | conserved Plasmodium protein unknown function | 60 | 1137.80 | | 2.28 | |
| | PF3D7_0914700 | major facilitator superfamily-related transporter putative | 50 | 479.60 | | 0.96 | |
| | PF3D7_0919900 | regulator of chromosome condensation-PP1-interacting protein | 60 | 620.45 | | 1.24 | |
| | PF3D7_1014900 | conserved Plasmodium protein unknown function | 60 | 1221.07 | | 2.44 | |
| | PF3D7_1031600 | conserved Plasmodium protein unknown function | 60 | 19500.89 | | 39.00 | |
| | PF3D7_1037300 | ADP/ATP transporter on adenylate translocase | 50 | 240.00 | | 0.48 | |
| | PF3D7_1109900 | 60S ribosomal protein L36 | 50 | 150.27 | | 0.30 | |
| | PF3D7_1116800 | heat shock protein 101 | 50 | 817.34 | | 1.63 | |
| | PF3D7_1121600 | exported protein 1 | 50 | 982.29 | | 1.96 | |
| | PF3D7_1124700 | GrpE protein homolog mitochondrial putative | 60 | 735.92 | | 1.47 | |
| | PF3D7_1129000 | spermidine synthase | 50 | 265.77 | | 0.53 | |
| | PF3D7_1134000 | heat shock protein 70 | 60 | 994.72 | | 1.99 | |
| | PF3D7_1211900 | non-SERCA-type Ca2+ -transporting P-ATPase | 50 | 430.33 | | 0.86 | |
| | PF3D7_1341300 | 60S ribosomal protein L18-2 putative | 50 | 534.94 | | 1.07 | |
| | PF3D7_1352800 | vacuolar fusion protein MON1 putative | 60 | 384.28 | | 0.77 | |
| | PF3D7_1360800 | falcilysin | 60 | 749.91 | | 1.50 | |
| | PF3D7_1417500 | H/ACA ribonucleoprotein complex subunit 4 putative | 60 | 784.24 | | 1.57 | |
| | PF3D7_1431700 | 60S ribosomal protein L14 putative | 50 | 665.11 | | 1.33 | |
| | PF3D7_1434800 | mitochondrial acidic protein MAM33 putative | 60 | 267.45 | | 0.53 | |
| | PF3D7_1471100 | exported protein 2 | 50 | 1188.97 | | 2.38 | |
| **MMV665806 50°C; N=985 proteins 55°C; N=851 proteins 60°C; N=676 proteins** | PF3D7_1211900 | non-SERCA-type Ca2+ -transporting P-ATPase | 55 | 9885.98 | 400.00 | 24.71 | 8.35 |
| | PF3D7_1360800 | falcilysin | 55 | 1893.27 | | 4.73 | |
| | PF3D7_0819600 | conserved Plasmodium protein unknown function | 60 | 5817.26 | | 14.54 | |
| | PF3D7_0904600 | ubiquitin specific protease putative | 60 | 3247.26 | | 8.12 | |
| | PF3D7_1015200 | cysteine--tRNA ligase | 60 | 3398.16 | | 8.50 | |
| | PF3D7_1336800 | nuclear movement protein putative | 60 | 3279.47 | | 8.20 | |
| **Piperaquine 51°C; N=718 proteins 57°C; N=618 proteins** | PF3D7_1360800 | falcilysin | 57 | 79.82 | 40.00 | 2.00 | 2.00 |
| **Pyronaridine 50°C; N=875 proteins 55°C; N=747 proteins 60°C; N=593 proteins** | PF3D7_0102900 | aspartate--tRNA ligase | 60 | 39.75 | 30.00 | 1.33 | 1.31 |
| | PF3D7_0105800 | cyclin-dependent kinases regulatory subunit putative | 50 | 18.31 | | 0.61 | |
| | PF3D7_0113000 | glutamic acid-rich protein | 50 | 1534.27 | | 51.14 | |
| | PF3D7_0113000 | glutamic acid-rich protein | 55 | 403.34 | | 13.44 | |
| | PF3D7_0113000 | glutamic acid-rich protein | 60 | 596.09 | | 19.87 | |
| | PF3D7_0213100 | protein SIS1 | 55 | 30.63 | | 1.02 | |
| | PF3D7_0213100 | protein SIS1 | 60 | 30.87 | | 1.03 | |
| | PF3D7_0305200 | conserved Plasmodium protein unknown function | 50 | 60.35 | | 2.01 | |
| | PF3D7_0305600 | DNA-(apurinic or apyrimidinic site) lyase putative | 55 | 56.93 | | 1.90 | |
| | PF3D7_0311300 | phosphatidylinositol 3- and 4-kinase putative | 55 | 62.31 | | 2.08 | |

| | | | | |
|---|---|---|---|---|
| PF3D7_0311300 | phosphatidylinositol 3- and 4-kinase putative | 60 | 28.73 | 0.96 |
| PF3D7_0505500 | DNA mismatch repair protein MSH6 putative | 50 | 38.48 | 1.28 |
| PF3D7_0505900 | mediator of RNA polymerase II transcription subunit 11 putative | 60 | 40.08 | 1.34 |
| PF3D7_0509100 | structural maintenance of chromosomes protein 4 putative | 50 | 14.05 | 0.47 |
| PF3D7_0513800 | ras-related protein Rab-1A | 55 | 54.45 | 1.82 |
| PF3D7_0520200 | mediator of RNA polymerase II transcription subunit 17 putative | 55 | 72.73 | 2.42 |
| PF3D7_0602100 | ATP-dependent RNA helicase putative | 50 | 92.86 | 3.10 |
| PF3D7_0603400 | trophozoite exported protein 1 | 50 | 21.50 | 0.72 |
| PF3D7_0603400 | trophozoite exported protein 1 | 60 | 39.45 | 1.31 |
| PF3D7_0604100 | AP2 domain transcription factor | 50 | 21.96 | 0.73 |
| PF3D7_0622800 | leucine--tRNA ligase putative | 55 | 37.08 | 1.24 |
| PF3D7_0622800 | leucine--tRNA ligase putative | 60 | 12.28 | 0.41 |
| PF3D7_0628100 | HECT-domain (ubiquitin-transferase) putative | 55 | 38.99 | 1.30 |
| PF3D7_0714500 | transcription elongation factor s-II putative | 50 | 25.66 | 0.86 |
| PF3D7_0714500 | transcription elongation factor s-II putative | 55 | 54.34 | 1.81 |
| PF3D7_0717200 | conserved Plasmodium protein unknown function | 55 | 36.96 | 1.23 |
| PF3D7_0717700 | serine--tRNA ligase putative | 60 | 20.77 | 0.69 |
| PF3D7_0804600 | tRNA pseudouridine synthase putative | 55 | 47.60 | 1.59 |
| PF3D7_0810300 | protein phosphatase PPM5 putative | 60 | 36.40 | 1.21 |
| PF3D7_0815600 | eukaryotic translation initiation factor 3 subunit G putative | 50 | 40.69 | 1.36 |
| PF3D7_0815600 | eukaryotic translation initiation factor 3 subunit G putative | 55 | 24.66 | 0.82 |
| PF3D7_0826100 | HECT-like E3 ubiquitin ligase putative | 50 | 68.91 | 2.30 |
| PF3D7_0826100 | HECT-like E3 ubiquitin ligase putative | 60 | 96.59 | 3.22 |
| PF3D7_0826500 | ubiquitin conjugation factor E4 B putative | 55 | 25.12 | 0.84 |
| PF3D7_0826500 | ubiquitin conjugation factor E4 B putative | 60 | 10.05 | 0.33 |
| PF3D7_0904000 | GTPase-activating protein putative | 60 | 36.18 | 1.21 |
| PF3D7_0913200 | elongation factor 1-beta | 50 | 45.24 | 1.51 |
| PF3D7_0913200 | elongation factor 1-beta | 60 | 34.55 | 1.15 |
| PF3D7_0921900 | conserved Plasmodium protein unknown function | 60 | 141.71 | 4.72 |
| PF3D7_0923900 | polyadenylate-binding protein 2 putative | 50 | 101.73 | 3.39 |
| PF3D7_0923900 | polyadenylate-binding protein 2 putative | 60 | 26.67 | 0.89 |
| PF3D7_1004300 | E3 ubiquitin-protein ligase putative | 50 | 9.30 | 0.31 |
| PF3D7_1004300 | E3 ubiquitin-protein ligase putative | 55 | 34.07 | 1.14 |
| PF3D7_1004300 | E3 ubiquitin-protein ligase putative | 60 | 25.24 | 0.84 |
| PF3D7_1010600 | eukaryotic translation initiation factor 2 subunit beta | 50 | 42.84 | 1.43 |
| PF3D7_1010600 | eukaryotic translation initiation factor 2 subunit beta | 55 | 33.75 | 1.12 |
| PF3D7_1010600 | eukaryotic translation initiation factor 2 subunit beta | 60 | 32.98 | 1.10 |
| PF3D7_1015200 | cysteine--tRNA ligase | 60 | 20.24 | 0.67 |
| PF3D7_1015300 | methionine aminopeptidase 1b putative | 60 | 3628.24 | 120.94 |
| PF3D7_1033100 | S-adenosylmethionine decarboxylase/ornithine decarboxylase | 60 | 516.46 | 17.22 |

| | | | | |
|---|---|---|---|---|
| PF3D7_1118000 | conserved Plasmodium protein unknown function | 50 | 928.76 | 30.96 |
| PF3D7_1118300 | insulinase putative | 55 | 87.57 | 2.92 |
| PF3D7_1123100 | calcium-dependent protein kinase 7 | 55 | 41.07 | 1.37 |
| PF3D7_1128100 | prefoldin subunit 5 putative | 60 | 20.91 | 0.70 |
| PF3D7_1132300 | nucleic acid binding protein putative | 60 | 32.20 | 1.07 |
| PF3D7_1133800 | RNA (uracil-5-)methyltransferase putative | 55 | 16.00 | 0.53 |
| PF3D7_1133800 | RNA (uracil-5-)methyltransferase putative | 60 | 10.20 | 0.34 |
| PF3D7_1138500 | protein phosphatase PPM2 | 50 | 33.99 | 1.13 |
| PF3D7_1138500 | protein phosphatase PPM2 | 55 | 50.98 | 1.70 |
| PF3D7_1138500 | protein phosphatase PPM2 | 60 | 54.91 | 1.83 |
| PF3D7_1141700 | OTU domain-containing protein putative | 50 | 108.50 | 3.62 |
| PF3D7_1141700 | OTU domain-containing protein putative | 55 | 67.08 | 2.24 |
| PF3D7_1206200 | eukaryotic translation initiation factor 3 subunit C putative | 50 | 34.80 | 1.16 |
| PF3D7_1206200 | eukaryotic translation initiation factor 3 subunit C putative | 60 | 38.15 | 1.27 |
| PF3D7_1212700 | eukaryotic translation initiation factor 3 subunit A putative | 50 | 37.06 | 1.24 |
| PF3D7_1212700 | eukaryotic translation initiation factor 3 subunit A putative | 55 | 32.53 | 1.08 |
| PF3D7_1212700 | eukaryotic translation initiation factor 3 subunit A putative | 60 | 33.69 | 1.12 |
| PF3D7_1213800 | proline--tRNA ligase | 60 | 4.80 | 0.16 |
| PF3D7_1216900 | DNA-binding chaperone putative | 55 | 21.25 | 0.71 |
| PF3D7_1224300 | polyadenylate-binding protein 1 putative | 50 | 80.87 | 2.70 |
| PF3D7_1233600 | asparagine and aspartate rich protein 1 | 55 | 27.87 | 0.93 |
| PF3D7_1244100 | N-alpha-acetyltransferase 15 NatA auxiliary subunit putative | 60 | 39.50 | 1.32 |
| PF3D7_1304500 | small heat shock protein putative | 50 | 100.56 | 3.35 |
| PF3D7_1304500 | small heat shock protein putative | 55 | 112.50 | 3.75 |
| PF3D7_1304500 | small heat shock protein putative | 60 | 197.73 | 6.59 |
| PF3D7_1308200 | carbamoyl phosphate synthetase | 60 | 619.49 | 20.65 |
| PF3D7_1319400 | conserved Plasmodium protein unknown function | 50 | 8.43 | 0.28 |
| PF3D7_1331700 | glutamine--tRNA ligase putative | 60 | 59.89 | 2.00 |
| PF3D7_1349200 | glutamate--tRNA ligase putative | 55 | 61.07 | 2.04 |
| PF3D7_1355500 | serine/threonine protein phosphatase 5 | 60 | 34.94 | 1.16 |
| PF3D7_1359400 | CUGBP Elav-like family member 1 | 60 | 34.42 | 1.15 |
| PF3D7_1360800 | falcilysin | 60 | 22.20 | 0.74 |
| PF3D7_1360900 | RNA-binding protein putative | 50 | 49.55 | 1.65 |
| PF3D7_1401800 | choline kinase | 55 | 36.99 | 1.23 |
| PF3D7_1412100 | mini-chromosome maintenance complex-binding protein putative | 60 | 32.13 | 1.07 |
| PF3D7_1417800 | DNA replication licensing factor MCM2 | 55 | 51.45 | 1.72 |
| PF3D7_1417800 | DNA replication licensing factor MCM2 | 60 | 43.47 | 1.45 |
| PF3D7_1434300 | Hsp70/Hsp90 organizing protein | 50 | 49.36 | 1.65 |
| PF3D7_1435500 | clathrin light chain putative | 50 | 33.98 | 1.13 |
| PF3D7_1442300 | tRNA import protein tRIP | 55 | 57.35 | 1.91 |
| PF3D7_1442300 | tRNA import protein tRIP | 60 | 37.81 | 1.26 |

| | | | | |
|---|---|---|---|---|
| PF3D7_1442900 | protein transport protein SEC7 putative | 50 | 40.78 | 1.36 |
| PF3D7_1442900 | protein transport protein SEC7 putative | 55 | 40.38 | 1.35 |
| PF3D7_1457300 | MA3 domain-containing protein putative | 50 | 41.01 | 1.37 |
| PF3D7_1457300 | MA3 domain-containing protein putative | 55 | 61.16 | 2.04 |
| PF3D7_1457300 | MA3 domain-containing protein putative | 60 | 34.17 | 1.14 |
| PF3D7_1473200 | DnaJ protein putative | 50 | 55.64 | 1.85 |
| PF3D7_1473200 | DnaJ protein putative | 55 | 52.59 | 1.75 |
| PF3D7_1473200 | DnaJ protein putative | 60 | 39.65 | 1.32 |

### Table S2. Proteins Identified as Protein Hits in Lysate CETSA

| Drug | Gene ID | Description | Heat challenge (°C) | MDT (nM) | *In vitro* $IC_{50}$ (nM) | $MDT/IC_{50}$ ratio | Median MDT/IC50 ratio |
|---|---|---|---|---|---|---|---|
| **Amodiaquine 51°C; N=1685 proteins 57°C; N=1626 proteins** | PF3D7_0213500 | tetratricopeptide repeat protein putative | 51 | 546.76 | 15.00 | 36.45 | 2257.30 |
| | PF3D7_0513300 | purine nucleoside phosphorylase | 57 | 33859.47 | | 2257.30 | |
| | PF3D7_1118000 | conserved Plasmodium protein unknown function | 51 | 21947.90 | | 1463.19 | |
| | PF3D7_1118000 | conserved Plasmodium protein unknown function | 57 | 59854.56 | | 3990.30 | |
| | PF3D7_1136500 | casein kinase 1 | 51 | 69803.64 | | 4653.58 | |
| **Chloroquine 51°C; N=1735 proteins 57°C; N=1540 proteins** | PF3D7_0107800 | double-strand break repair protein MRE11 | 57 | 422.90 | 25.00 | 16.92 | 927.11 |
| | PF3D7_0308000 | DNA polymerase delta small subunit putative | 57 | 88400.24 | | 3536.01 | |
| | PF3D7_0619400 | cell division cycle protein 48 homologue putative | 57 | 35656.94 | | 1426.28 | |
| | PF3D7_0812600 | ubiquitin-conjugating enzyme E2 putative | 51 | 304.08 | | 12.16 | |
| | PF3D7_1118000 | conserved Plasmodium protein unknown function | 51 | 6323.59 | | 252.94 | |
| | PF3D7_1118000 | conserved Plasmodium protein unknown function | 57 | 91918.12 | | 3676.72 | |
| | PF3D7_1142500 | 60S ribosomal protein L28 | 51 | 64742.79 | | 2589.71 | |
| | PF3D7_1315400 | zinc finger (CCCH type) protein putative | 57 | 3158.75 | | 126.35 | |
| | PF3D7_1329400 | AMP deaminase putative | 57 | 79.97 | | 3.20 | |
| | PF3D7_1360800 | falcilysin | 57 | 56124.73 | | 2244.99 | |
| | PF3D7_1424400 | 60S ribosomal protein L7-3 putative | 51 | 196158.95 | | 7846.36 | |
| | PF3D7_1434800 | mitochondrial acidic protein MAM33 putative | 57 | 10698.59 | | 427.94 | |
| **Lumefantrine 51°C; N=2055 proteins 57°C; N=2079 proteins** | PF3D7_0309300 | N2227-like protein putative | 51 | 39.51 | 20.00 | 1.98 | 169.14 |
| | PF3D7_1334300 | MSP7-like protein | 57 | 6726.14 | | 336.31 | |
| **MK-4815 51°C; N=1628 proteins 57°C; N=1528 proteins** | PF3D7_0317700 | CPSF (cleavage and polyadenylation specific factor) subunit A putative | 51 | 8013.10 | 250.00 | 32.05 | 7.44 |
| | PF3D7_0618300 | 60S ribosomal protein L27a putative | 57 | 3438.68 | | 13.75 | |
| | PF3D7_1105100 | histone H2B | 57 | 283.57 | | 1.13 | |
| | PF3D7_1124600 | ethanolamine kinase | 51 | 12599.03 | | 50.40 | |
| | PF3D7_1234900 | conserved Plasmodium protein unknown function | 51 | 203.06 | | 0.81 | |
| | PF3D7_1360800 | falcilysin | 57 | 87.26 | | 0.35 | |
| **MMV000848 50°C; N=1906 proteins 55°C; N=1787 proteins 60°C; N=1676 proteins** | PF3D7_0210100 | 60S ribosomal protein L37ae putative | 55 | 2503.25 | 500.00 | 5.01 | 0.96 |
| | PF3D7_0513300 | purine nucleoside phosphorylase | 60 | 242.58 | | 0.49 | |
| | PF3D7_0513300 | purine nucleoside phosphorylase | 55 | 104.72 | | 0.21 | |
| | PF3D7_0531100 | conserved Plasmodium protein unknown function | 55 | 479.89 | | 0.96 | |
| | PF3D7_1358800 | 40S ribosomal protein S15 | 60 | 4837.39 | | 9.67 | |
| | PF3D7_1360800 | falcilysin | 60 | 1108.95 | | 2.22 | |
| | PF3D7_1416500 | NADP-specific glutamate dehydrogenase | 60 | 2.98 | | 0.01 | |

| | | | | | | | |
|---|---|---|---|---|---|---|---|
| **MMV665806 50°C; N=1851 proteins 55°C; N=1686 proteins 60°C; N=1579 proteins** | PF3D7_1360800 | falcilysin | 60 | 1026.45 | 400.00 | 2.57 | 2.57 |
| **Piperaquine 51°C; N=2017 proteins 57°C; N=1685 proteins** | PF3D7_0502900 | Tim10/DDP family zinc finger protein putative | 57 | 1074.97 | 10.00 | 107.50 | 0.61 |
| | PF3D7_0532400 | lysine-rich membrane-associated PHISTb protein | 57 | 6.13 | | 0.61 | |
| | PF3D7_1248200 | pre-mRNA-splicing factor RBM22 putative | 57 | 0.10 | | 0.01 | |
| **Pyronaridine 51°C; N=2085 proteins 57°C; N=2036 proteins** | PF3D7_0313100 | ubiquitin-protein ligase putative | 57 | 36.40 | 5.00 | 7.28 | 7.28 |
| | PF3D7_0313400 | conserved Plasmodium protein unknown function | 57 | 2.74 | | 0.55 | |
| | PF3D7_1323400 | 60S ribosomal protein L23 | 57 | 32491.81 | | 6498.36 | |
| | PF3D7_1234900 | conserved Plasmodium protein unknown function | 51 | 30483.85 | | 6096.77 | |
| | PF3D7_1409200 | conserved Plasmodium protein unknown function | 57 | 20.49 | | 4.10 | |

**Table S3. Crytallography Refinement Parameters**

| | CQ | MFQ | MK-4815 | MMV665806 | MMV000848 |
|---|---|---|---|---|---|
| **Crystallisation condition** | 0.2 M sodium acetate trihydrate pH 8.0, 20% w/v PEG 3,350 | 0.2 M sodium acetate trihydrate pH 7.0, 20% w/v PEG 3,350 | 0.2 M ammonium acetate, 0.1 M Tris pH 8.5, 25% w/v PEG 3,350 | Morpheus C9<br><br>(pH 8.5) | Morpheus C5<br><br>(pH 7.5) |
| **PDB accession** | 7DI7 | 7DIA | 7DIJ | 8HO5 | 8HO4 |
| **Wavelength (Å)** | 1 | 1 | 1 | 1 | 1 |
| **Resolution range (Å)** | 49.2 - 1.821 (1.887 - 1.821)[a] | 46.91 - 1.551 (1.606 - 1.551) | 46.84 - 1.9 (1.968 - 1.9) | 47.12 - 2.003 (2.074 - 2.003) | 47.4 - 1.96 (2.03 - 1.96) |
| **Space group** | $P2_12_12_1$ | $P2_12_12_1$ | $P2_12_12_1$ | $P2_12_12_1$ | $P2_12_12_1$ |
| **Unit cell (Å, °)** | 93.94 106.61 127.81 90 90 90 | 93.46 105.74 126.35 90 90 90 | 93.52 105.83 125.93 90 90 90 | 93.89 106.29 126.91 90 90 90 | 94.26 106.61 127.89 90 90 90 |
| **Total reflections** | 1555036 (154468) | 2424712 (233346) | 1308423 (127012) | 839013<br>-80956 | 1254171 (111073) |
| **Unique reflections** | 115028 (11325) | 180859 (17886) | 98799 (9661) | 85579 (8370) | 92720 (8971) |
| **Multiplicity** | 13.5 (13.6) | 13.4 (13.0) | 13.2 (13.1) | 9.8 (9.7) | 13.5 (12.4) |
| **Completeness (%)** | 99.94 (99.61) | 99.96 (99.68) | 99.85 (98.83) | 99.56 (98.81) | 99.72 (97.83) |
| **Mean I/sigma(I)** | 16.88 (1.94) | 23.27 (1.81) | 15.48 (1.85) | 14.65 (2.38) | 28.00 (4.92) |
| **Wilson B-factor** | 21.75 | 20.01 | 26.65 | 32.63 | 29.7 |
| **$R_{merge}$[b]** | 0.1393 (1.347) | 0.07683 (1.397) | 0.1433 (1.358) | 0.1257 (1.252) | 0.0615 (0.501) |
| **$R_{meas}$** | 0.1447 (1.399) | 0.07989 (1.454) | 0.1491 (1.413) | 0.1329 (1.322) | 0.06392 (0.5225) |
| **$R_{pim}$** | 0.03914 (0.3748) | 0.02176 (0.4002) | 0.04084 (0.3852) | 0.04227 (0.416) | 0.01727 (0.1464) |
| **CC1/2** | 0.999 (0.7) | 1 (0.706) | 0.999 (0.738) | 0.998 (0.757) | 1 (0.948) |
| **CC*** | 1 (0.907) | 1 (0.91) | 1 (0.922) | 0.999 (0.928) | 1 (0.986) |
| **Reflections used in refinement** | 115014 (11325) | 180850 (17884) | 98767 (9656) | 85571 (8370) | 92713 (8971) |
| **Reflections used for $R_{free}$** | 5750 (566) | 9043 (894) | 4938 (482) | 4277 (418) | 4636 (448) |
| **$R_{work}$[c]** | 0.1760 (0.2990) | 0.1741 (0.2831) | 0.1938 (0.3474) | 0.1778 (0.2921) | 0.1712 (0.2207) |
| **$R_{free}$[d]** | 0.1981 (0.3502) | 0.1972 (0.3057) | 0.2094 (0.3597) | 0.1906 (0.3153) | 0.2048 (0.2436) |
| **CC(work)** | 0.963 (0.827) | 0.964 (0.834) | 0.952 (0.753) | 0.959 (0.837) | 0.961 (0.895) |

| | | | | | |
|---|---|---|---|---|---|
| **CC(free)** | 0.960 (0.780) | 0.953 (0.840) | 0.944 (0.719) | 0.957 (0.797) | 0.943 (0.847) |
| **Number of non-hydrogen atoms** | 10418 | 10353 | 9680 | 9775 | 9633 |
| **Protein residues** | 1076 | 1072 | 1083 | 1075 | 1076 |
| **RMS(bonds)** | 0.007 | 0.007 | 0.009 | 0.013 | 0.008 |
| **RMS(angles)** | 1.14 | 1.15 | 1.23 | 1.13 | 0.96 |
| **Ramachandran favored (%)** | 97.66 | 98.21 | 97.47 | 98.31 | 98.31 |
| **Ramachandran allowed (%)** | 2.34 | 1.79 | 2.53 | 1.69 | 1.69 |
| **Ramachandran outliers (%)** | 0 | 0 | 0 | 0 | 0 |
| **Rotamer outliers (%)** | 0.8 | 0.6 | 0.2 | 0 | 0.8 |
| **Clashscore** | 3.98 | 5.04 | 3.71 | 4.82 | 3.59 |
| **Average B-factor** | 24.91 | 25.65 | 28.02 | 39.18 | 33.46 |
| **macromolecules** | 23.17 | 23.97 | 27.55 | 38.4 | 32.91 |
| **ligands** | 43.88 | 40.33 | 35.94 | 54.33 | 52.04 |
| **solvent** | 34.89 | 36.23 | 33.19 | 46.42 | 39.49 |

[a] Statistics for the highest-resolution shell are shown in parentheses.

[b] $R_{merge} = \Sigma_h \Sigma_i |I_{hi} - \langle I_h \rangle| / \Sigma_{h,i} I_{hi}$, where $I_{hi}$ is the ith observation of the reflection h, while $\langle I_h \rangle$ is its mean intensity.

[c] R factor = $\Sigma ||F_{obs}| - |F_{calc}|| / \Sigma |F_{obs}|$.

[d] $R_{free}$ was calculated with 5% of reflections excluded from the whole refinement procedure.

**Table S4. List of Primers for Validation of FLN Transfectants**

| Primer name | Sequence (5`-3`) |
|---|---|
| P1 | GGTGACCATTCAGAGGAGAAGG |
| P2 | ACGTGATCTACCGCTAACGC |
| P3 | CTATGCGGCATCAGAGCAGA |
| P4 | ACGTGATCTACCGCTAACGC |
| P5 | TGGTGACCATTCAGAGGAGA |
| P6 | ACAAGGTGTATAATGTAAGTTTGCT |
| P7 | GTCGCTGTGTCCCAGAACAT |
| P8 | ATGGCCTGGGTGATTCATGG |
| P_OE1 | TAAGCACCATGGATGAATTTAACAAAATTAAT |
| P_OE2 | TGCTTAGCTAGCTTCTATTAATACCTTTTTA |
